# A Multiomic Analysis of Cachectic Mice Reveals Cancer Driven Suppression of Muscle Stem Cell Differentiation

**DOI:** 10.64898/2026.01.12.699072

**Authors:** Darren M. Blackburn, Aldo Hernández-Corchado, Korin Sahinyan, Hedieh Habibi Khorasani, Felicia Lazure, Vincent Richard, Dianbo Qu, Shuo Wang, Christoph H. Borchers, Arezu Jahani-Asl, Hamed S. Najafabadi, Antonis E. Koromilas, Vahab D. Soleimani

**Affiliations:** Department of Biochemistry, Microbiology & Immunology, University of Ottawa, 451 Smyth Rd, Ottawa, ON, K1H 8M5 Canada; Department of Human Genetics, McGill University, 3640 rue University, Montréal, QC, H3A 0C7, Canada; Lady Davis Institute for Medical Research, Jewish General Hospital, 3755 Chemin de la Côte-Sainte-Catherine, Montréal QC, H3T 1E2, Canada; Moffit Cancer Center, Tumor Microenvironment and Metastasis Department, Tampa, FL 33612, USA; Segal Cancer Proteomics Centre, Lady Davis Institute, Jewish General Hospital, McGill University, Montréal, Quebec H3T 1E2, Canada; Gerald Bronfman Department of Oncology, Lady Davis Institute for Medical Research, Jewish General Hospital, Montréal, QC H3T 1E2, Canada; Division of Experimental Medicine, McGill University, Montréal, QC H4A 3J1, Canada; Department of Pathology, McGill University, Montréal, QC H3A 2B4, Canada; Department of Cellular and Molecular Medicine and University of Ottawa Brain and Mind Research Institute, University of Ottawa, Ottawa, ON, Canada

## Abstract

Cancer cachexia affects a large proportion of cancer patients, inducing a rapid decline in muscle mass. Patients with cachexia have a worse prognosis and are less responsive to cancer therapies. The exact cause of cachexia remains unknown, nor are there any effective treatments for the condition. In this study, we use the C26 adenocarcinoma cell line to determine how cancer cells affect myofiber and muscle stem cell function. We determined that C26 cancer cells adapt to the host environment, in both male and female mice, greatly altering their transcriptome to promote their survival and growth. C26 cells directly communicate with muscle stem cells via GDF15 and MMP9. These circulatory factors cause the muscle stem cells to upregulate the EMT pathway and become less capable of undergoing differentiation and contributing to muscle regeneration. Muscle stem cells from tumor bearing mice are less proliferative and less prone to differentiation, Chromatin accessibility data shows that there are fewer accessible myogenic regulatory binding sites. Cytokine array determined that circulating GDF15 and MMP9 were highly upregulated and were derived form C26 tumor cells. However, blocking tumor derived GDF15 is not sufficient to prevent the onset of cachexia and rescue the loss of muscle stem cell function. Together, these findings establish a new conceptual paradigm in which cancer orchestrates muscle wasting through coordinated transcriptional, metabolic, and epigenetic suppression of muscle stem cell differentiation.

## Introduction

Cancer cachexia is a debilitating condition where patients suffer a severe and rapid decline in muscle mass, with or without the loss of fat mass^1-3^. Muscle wasting is a common symptom in cancer, with approximately 80% of late-stage cancer patients exhibiting it^4^. Cachexia results in a worse prognosis for patients; they are less responsive to cancer therapies and suffer a loss of quality of life^1,5,6^. Furthermore, cachexia is itself responsible for approximately 30% of all cancer related death, primarily due to respiratory or heart failure^4^. Beyond muscle wasting, cancer cachexia is also associated with metabolic imbalance, systemic inflammation, insulin resistance, and anorexia^7-10^. It is a complex and multifactorial condition.

Cancer cachexia is not present in all cancers equally. It is more prevalent in gastrointestinal and pancreatic cancers, with between 60-70% of patients exhibiting muscle wasting^11^. However, it is much less common in breast cancer, with muscle wasting affecting nearly 20% of patients^11^. It is important to note, that the cancer does not need to be present in the muscle to induce muscle wasting. Other factors that affect the incidence of cachexia are the stage of the cancer; patients with advanced cancer are more likely to have cancer cachexia compared to those in earlier stages, as well as the sex of the patient^12^. Males are more susceptible to cancer cachexia, present in 61% of male patients in contrast to the 31% incidence in female patients^12,13^.

Despite the severity and prevalence of the condition, there is no effective treatment or therapy for cachexia. Furthering our understanding on how the cancer cells alter skeletal muscle and muscle stem cell behaviour is necessary for the development of effective therapies. In this study, we sequenced the transcriptome of myofibers and muscle stem cells from healthy controls and tumor bearing male and female mice, identifying several key pathways that are deregulated. Muscle stem cells in cachectic mice exhibit delayed activation and a decrease in myogenic potential. Additionally, we discovered how C26 cancer cells alter their transcriptome to adapt to the host environment. Cachexia is a debilitating condition in which the molecular mechanisms driving it are not well understood. Here, we conduct a comprehensive overview of the changes occurring in the skeletal muscle, and the resident MuSCs.

## Results

### C26 Tumors Induce Cachexia in Male and Female Mice

We used C26 adenocarcinoma cells, a highly aggressive cancer that is known to induce cachexia^14,15^. 3-month-old male and female Balb/c mice were given two subcutaneous injections of C26 cells at different locations of the body, and the tumor were allowed to grow until the mice reached endpoint, typically between the 2 and 3 week postinjection timepoint, and the muscle was collected for analysis (Fig. 1a).

**Figure 1:**
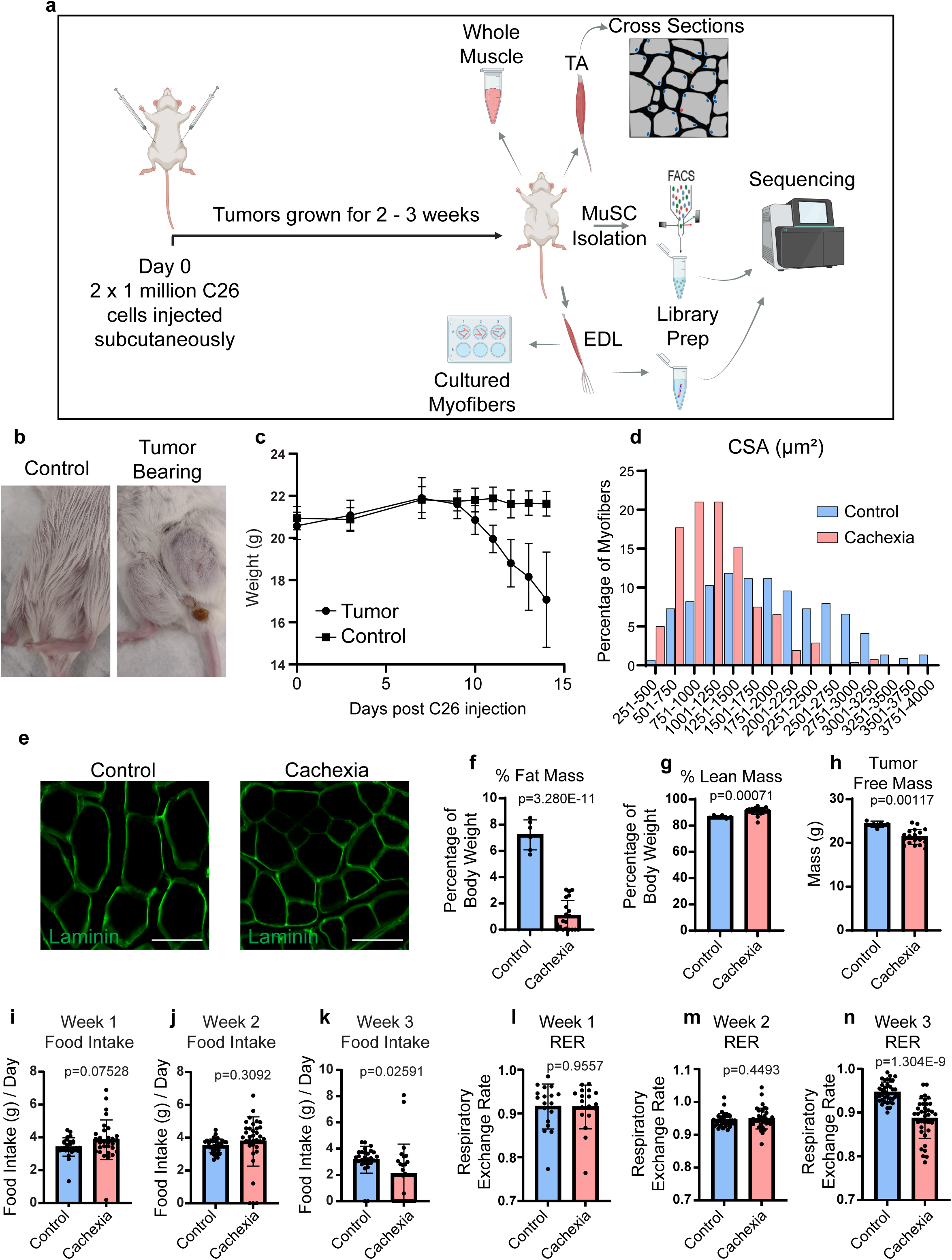
C26 tumors induce cachexia in male and female mice. **a.** Schematic of the overall experimental design for C26 tumor growth and muscle analysis. **b.** Image of female healthy control and tumor bearing Balb/c mice. **c.** Line chart of the change in body mass in female mice as tumor develops. n = 5 mice. **d.** Binning of the CSA of myofibers from healthy control and tumor bearing female mice. Data is presented as a percentage of all myofibers that measure within the marked range in area. n = 5 mice. **e.** Representative image of a TA cross section, scale bar = 70µm. **f.** Percentage of fat mass in male healthy control and cachectic mice, measured at endpoint (16-19 days post C26 injection). n = 6 controls, and 20 tumor bearing mice. **g.** Percentage of lean mass in male healthy control and cachectic mice, measured at endpoint (16-19 days post C26 injection). n = 6 controls, and 20 tumor bearing mice. **h.** The body weight of male mice after removal of tumors. n = 6 controls, and 20 tumor bearing mice. Daily measured food consumption in male mice in the **i.** first week, **j.** second week, and **k.** third week post tumor cell injection. n = 6 mice. Average daily RER value quantified in male mice in the **l.** first week, **m.** second week, and **n.** third week post C26 injection. n = 6 mice.

To begin, we wanted to validate the findings that the C26 cells do indeed cause cachexia. We see that the C26 cells can quickly form large tumors in the mice (Fig. 1b) and that the female mice show a rapid and large loss in body weight (Fig. 1c), as well as a decrease in myofiber size (Fig. 1d,e). We analyzed male mice using an EchoMRI at their endpoint and found a dramatic decrease in the percentage of fat mass, with many of the mice reaching 0% (Fig. 1f). As a consequence of this loss of fat, there is a subsequent increase in the percentage of lean mass in the tumor bearing mice (Fig. 1g). This may seem contrary to what cachexia is, which is the loss of muscle mass with or without the loss of fat, but because these are percentages the loss of fat will cause the percentage of the lean mass to increase. It is also important to note that the tumor itself is measured primarily as lean mass in the EchoMRI (Data not shown). As a result, we see a large decrease in the tumor free body mass of these male mice (Fig. 1h).

Cachexia is associated with changes in diet and metabolism^16,17^. To quantify these changes, we housed male mice in metabolic cages for the entire period of tumor development, from tumor injection to endpoint. We see that within the first 2 weeks of tumor growth that there is no significant difference between the healthy controls and the tumor bearing mice with regards to food consumption (Fig. 1i,j). However, within the third week, we see a significant drop in the amount of food eaten by the tumor bearing mice (Fig. 1k). This is reflected in the energy metabolism as well. Within the first 2 weeks, there is no difference in the metabolism, with both conditions closer to a Respiratory Exchange Ratio (RER) of 1.0, indicating a preferential use of carbohydrates for their energy (Fig. 1l,m). Then in the third week, RER values in the cachectic mice drops towards 0.7 (Fig. 1n), indicating that they are starting to use more fat for energy, which may explain the loss of fat mass in these mice. Overall, we confirmed that the C26 cells that will be used in the rest of this study do induce cachexia.

### Cancer Alters the Myofiber Transcriptome

Myofibers are the primary component of skeletal muscle, and they undergo atrophy in cachexia. However, to date, there has been no sequencing performed on only the myofibers to determine the transcriptomic changes that cachexia induces. To address this knowledge gap, we performed single myofiber RNA-Seq (smfRNA-Seq)^18^ on individual myofibers isolated from healthy controls or tumor bearing mice, in both male and females (Fig. 2a). From the RNA-Seq results, we see that there is a large change in the myofiber transcriptome due to cancer, in both male and female mice (Fig. 2b,c). Using a threshold of LFC ≥ 1, adjusted p-value ≤ 0.05 and RPM ≥ 5 in at least one condition, we see in female myofibers that 770 genes are significantly downregulated, and 1154 genes are upregulated when comparing myofibers from healthy controls to those from tumor bearing mice (Fig. 2d,e). This is in contrast to male myofibers, where only 150 and 231 genes are significantly downregulated or upregulated, respectively (Fig. 2d,e).

**Figure 2:**
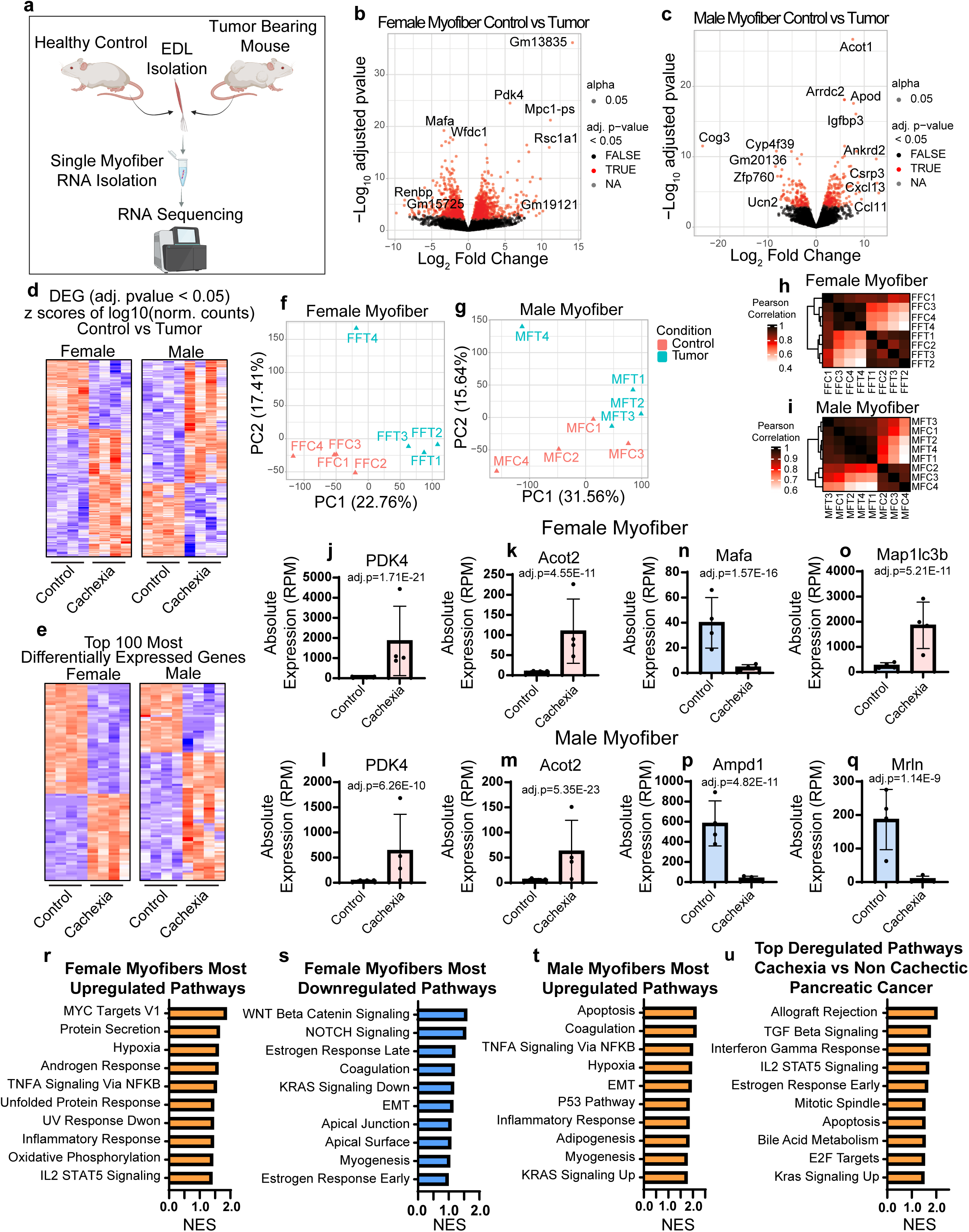
Cancer alters the myofiber transcriptome. **a.** Schematic of the experimental design, where the RNA from individual myofibers isolated from tumor bearing mice and healthy controls is used to generate single myofiber RNA-Seq libraries. **b.** Volcano plot of sequenced female myofibers from control and tumor bearing mice, genes are labelled as significantly different if the adj. pvalue ≤ 0.05. **c.** Volcano plot of sequenced male myofibers from control and tumor bearing mice, genes are labelled as significantly different if the adj. pvalue ≤ 0.05. **d.** Heatmaps of the differentially expressed genes (adj. pvalue ≤ 0.05) of control vs cachexia myofiber samples from both female and male mice. **e.** Heatmaps of the top 100 differentially expressed genes (adj. pvalue ≤ 0.05) of control vs cachexia myofiber samples from both female and male mice. **f.** PCA plot of myofibers from control and tumor bearing female mice. **g.** PCA plot of myofibers from control and tumor bearing male mice. **h.** Pearson correlations between the sequenced female myofiber samples. **i.** Pearson correlations between the sequenced male myofiber samples. **j.** Bar graphs of the expression of PDK4 and **k.** Acot 2 in female myofibers. **l.** Bar graphs of the expression of PDK4 and **m.** Acot 2 in male myofibers. **n.** Mafa expression in female myofibers. **o.** Bar graph of the RPM value of Map1lc3b in female myofibers. **p.** RPM expression of Ampd1 in male myofibers. **q.** Mrln expression in male myofibers. **r.** Bar graph of the top ten most upregulated Hallmark pathways, in terms of enrichment score, from control vs tumor female myofibers. **s.** Bar graph of the top ten most downregulated Hallmark pathways, in terms of enrichment score, from control vs tumor female myofibers. **t.** Bar graph of the top ten most upregulated Hallmark pathways, in terms of enrichment score, from control vs tumor male myofibers. **u.** Bar graph of the top ten most significantly upregulated pathways from microarray data of skeletal muscle biopsies from cachectic vs non cachectic pancreatic ductal adenocarcinoma patients. Data was retrieved from GSE130563^14^.

The reason for this large discrepancy between the sexes is mostly due to the inherent variability between samples as each sequenced sample is that of an individual myofiber. We see that in PCA plots, the control and cachectic conditions do separate out from one another in both female and male samples (Fig. 2f,g) however in the pearson correlation heatmaps we do not see a strong division between control and cachexia samples (Fig. 2h). Because smfRNA-Seq is a single cell sequencing method, it is to be expected that there would be variation between samples. This variation would cause changes in gene expression to be less significant in terms of adjusted p-value, due to the spread of the samples. We see that if we remove the adjusted p-value and RPM cutoff from our threshold that the number of genes in which the LFC ≥ 1, when comparing healthy controls vs cachexia, are nearly equal between female and male myofibers. Males have 2571 downregulated genes and 2373 upregulated genes, while in contrast females have 2547 and 2576 genes that are down and upregulated respectively.

A more in-depth analysis of the transcriptomic changes reveals that many key genes for myofiber function are deregulated in cachexia. Here, we highlight some genes selected from the top 20 most differentially expressed genes as ranked by adjusted p-value. There is an overlap between male and female myofibers. For example, PDK4, ACOT1 and ACOT2 are all in the top 20 most differentially expressed genes in both male and females (Fig2. J-m). PDK4 inhibits pyruvate dehydrogenase, meaning it has a role glucose metabolism, and its overexpression may shift metabolism towards fatty acid oxidation^19^. ACOT1 and ACOT2 are genes that encode Acyl-CoA Hydrolase enzymes, which is important for fatty acid metabolism^20^. These findings indicate that there may be major changes in the metabolism of myofibers in cachexia. Female myofibers also showed a decrease in *Mafa* transcript, a key transcription factor in the maturation and maintenance of fast type myofibers (Fig 2. n)^21,22^. Male myofibers also have a significant decrease in *Mafa* expression, but it does not fall within the top 20 genes. Mafa is also known to promote the expression of Myh4^21^, and we see in both male and female myofibers that Myh4 expression is decreased (Data not shown). Lastly, for the female myofibers, we observed a large increase in the expression of *Map1lc3b,* a gene that has a known function in autophagy (Fig2. o)^23^. Looking at the top 20 most differentially expressed genes in male myofibers, we see that there is a near complete ablation of the expression of *Ampd1* (Fig. 2p). AMPD1 deficiency is a common mutation with a wide array of phenotypes, ranging from asymptomatic to exercise intolerance to muscle atrophy, however the more severe phenotypes cannot be attributed solely to mutations in the AMPD1 gene^24^. Interestingly, while *Ampd1* has lost almost all of its transcripts in males, in females there is only an approximate 40% loss of expression. We also see a considerable decrease in *Mrln* in male myofibers, but not in female myofibers (Fig. 2q). This gene has a function in regulating Ca^2+^ uptake in the sarcoplasmic reticulum by inhibiting SERCA and its deregulation could cause defects in the ability of the myofiber to contract properly^25^.

We performed a GSEA analysis between control and cachectic myofibers. We see in the females that among the most upregulated pathways there is Myc Targets V1, which are genes that are often activated as a response to cancer (Fig. 2r). There is also an enrichment in the Hypoxia pathway (Fig. 2r), which is known to induce muscle atrophy and inhibits myofiber growth^26^. When looking at the downregulated Hallmark pathways, we see that the 2 most enriched pathways are WNT Beta Catenin Signaling and NOTCH Signaling (Fig. 2s). Both of these signaling pathways are crucial for proper muscle function^27,28^. In males, many of the upregulated pathways are similar to the females, such as Hypoxia (Fig. 2t). However, their top pathway is Apoptosis (Fig. 2t), indicating that the myofibers may be undergoing cell death in cachexia. Interestingly, no downregulated pathways were significantly enriched in the male context. We wanted to compare how our results from mice contrasts with those from humans. A recent study by Judge et al. performed microarray on muscle biopsies from pancreatic ductal adenocarcinoma (PDAC) patients^14^. Using the publicly available data (GSE130563)^14^ we performed a GSEA between the cachectic (n=17) and non cachectic (n=5) patients. We found that there was a large overlap between the pathways that are enriched in the human cachectic patients and the ones enriched in the mouse data. Notably, IL2 STAT5 Signaling, Apoptosis and KRAS Signaling Up (Fig. 2u). TNFA Signaling via NFKB, IL6 JAK STAT3 Signaling and Inflammatory response were also enriched but did not reach significance. These results indicate that there are common pathways that are deregulated between our mouse and human data.

There are known sex differences in muscle function, therefore we wanted to further explore any potential differences between male and female myofibers and how they respond to cachexia. We see that there is a large difference in the transcriptome between male and female myofibers, regardless of whether they are from healthy mice or cachectic mice (Extended Data Fig. 1a). At a threshold of LFC ≥ 1, adjusted p-value ≤ 0.05 and RPM ≥ 5 in at least one condition, there are 1039 genes that are upregulated in males and 451 that are downregulated (Extended Data Fig. 1a) in healthy myofibers. In cachectic myofibers, there are 789 and 516 genes are that upregulated and downregulated in males (Extended Data Fig. 1a). Interestingly, the majority of the top 100 differentially expressed genes are upregulated in the males (Extended Data Fig. 1b), this may be partially due to the Y chromosome genes, which are uniquely expressed in male cells. Through PCA analysis and Pearson Correlations, the samples tend to cluster more based on sex, while still showing a high degree of variability (Extended Data Fig. 1c-f).

A more in-depth analysis of the genes that are differentially expressed between male and female myofibers indicates, as expected, that the majority of the top DEGs in are those coded on the Y chromosome, such as Ddx3y and Eif2s3y (Extended Data Fig. 1g,h). However, not every top DEG is on the Y chromosome. For example, we see that ZFP280d, which is present on Chromosome 9, is very highly expressed in male myofibers in healthy controls, averaging at 1842 RPM, while in females it is mostly not expressed with an average RPM of 2.7 (Extended Data Fig. 1g). Very little is known about ZFP280d, to date no studies have been performed on its role in muscle. Interestingly, in cachexia, males lose expression of ZFP280d, with the average expression dropping to 208 RPM (data not shown). Generally, we see that there is a positive correlation on the effect of cachexia on both male and females. When all genes are considered, there is an r value of 0.2432, a modest correlation. However, when we only plot genes that are significantly different in both comparisons, we see a much stronger correlation of r = 0.8435. As a whole, while there are differences on the effect of cachexia between male and female myofibers, the factor that more greatly explains their differences is sex itself.

An imbalance between protein synthesis and protein degradation is viewed to be the mechanism that causes the loss in muscle mass^29^. We wanted to assess the protein dynamics in whole muscle. To do so, we used heavy labelled amino acids to track the formation of new proteins during cachexia. In short, we treated Balb/c mice with heavy arginine in their drinking water for one week, after which the water was replaced with unsupplemented water. Two days later, the mice were injected with C26 cells and the tumors were allowed to grow for 7 days. Afterwards, mice were then given water that had heavy lysine in it for another 7 days. The mice were then euthanized and the whole muscle and tumors processed for Mass Spec analysis.

From our results, we see that there is a distinct change in the proteome of cachectic mice compared to healthy controls, with the samples clustering separately on the PCA (Extended Data Fig. 2a). Pearson analysis also indicates that the cachectic muscle samples are more similar to each other than the controls (Extended Data Fig. 2b). However, we only detected 29 significantly different (LFC ≥ 1.5, adjusted p-value ≤ 0.05) proteins, and interestingly, most of them were upregulated (Extended Data Fig. 2c,d). This low number of significantly different proteins may be due to challenges in performing mass spec in whole muscle as the structural proteins are very abundant. In our results, the top 20 most expressed proteins in the whole muscle, most of which are the myofiber structural genes, represent 50% of all detected peptides. Titin alone, which is the most highly expressed protein in the muscle, represents about 17% of all detected peptides in our results. This is in contrast to the whole tumor mass spec, where the top 20 proteins only represent about 18% of all detected peptides.

When quantifying the percentage of peptides containing the heavy amino acids, we see interesting results. To begin, the muscle from the control and the cachectic mice show the incorporation of an equal percentage of heavy amino acids. As expected, the first administered amino acid (arginine), was detected at lower levels and the second heavy amino acid was highly detected, indicating that the cachectic myofibers were still capable of incorporating the amino acid and therefore were producing new proteins (Extended Data Fig. 2e). However, the most interesting result from the Mass Spec data is that the tumor itself incorporated the heavy arginine, albeit at lower levels than the muscle, which was administered prior to tumor injection (Extended Data Fig. 2e). This suggests that the tumor cells are consuming the proteins from other tissues, most likely from the muscle as it is the protein storage of the body.

### Muscle Stem Cell Transcriptome is Greatly Altered in Cachexia

MuSCs are essential for muscle regeneration and maintenance. However, no study to date has analyzed the transcriptome of MuSCs in the context of cachexia. We have sequenced freshly isolated MuSCs from both male and female mice in 4-month-old healthy controls and tumor bearing mice (Fig.3 2a). From these results, we see that there is a large change in the transcriptome of MuSCs due to cachexia (Fig.3 2b,c). In female mice, at a threshold of LFC ≥ 1, adjusted p-value ≤ 0.05 and RPM ≥ 5 in at least one condition, MuSCs have 2185 genes down in cachexia and 2211 are upregulated (Fig.3 2d,e). Males in contrast have 751 and 531 genes that are downregulated and upregulated, respectively (Fig.3 2d,e). The MuSC samples cluster mostly based on cachexia in both males and females (Fig.3 2f-i).

**Figure 3:**
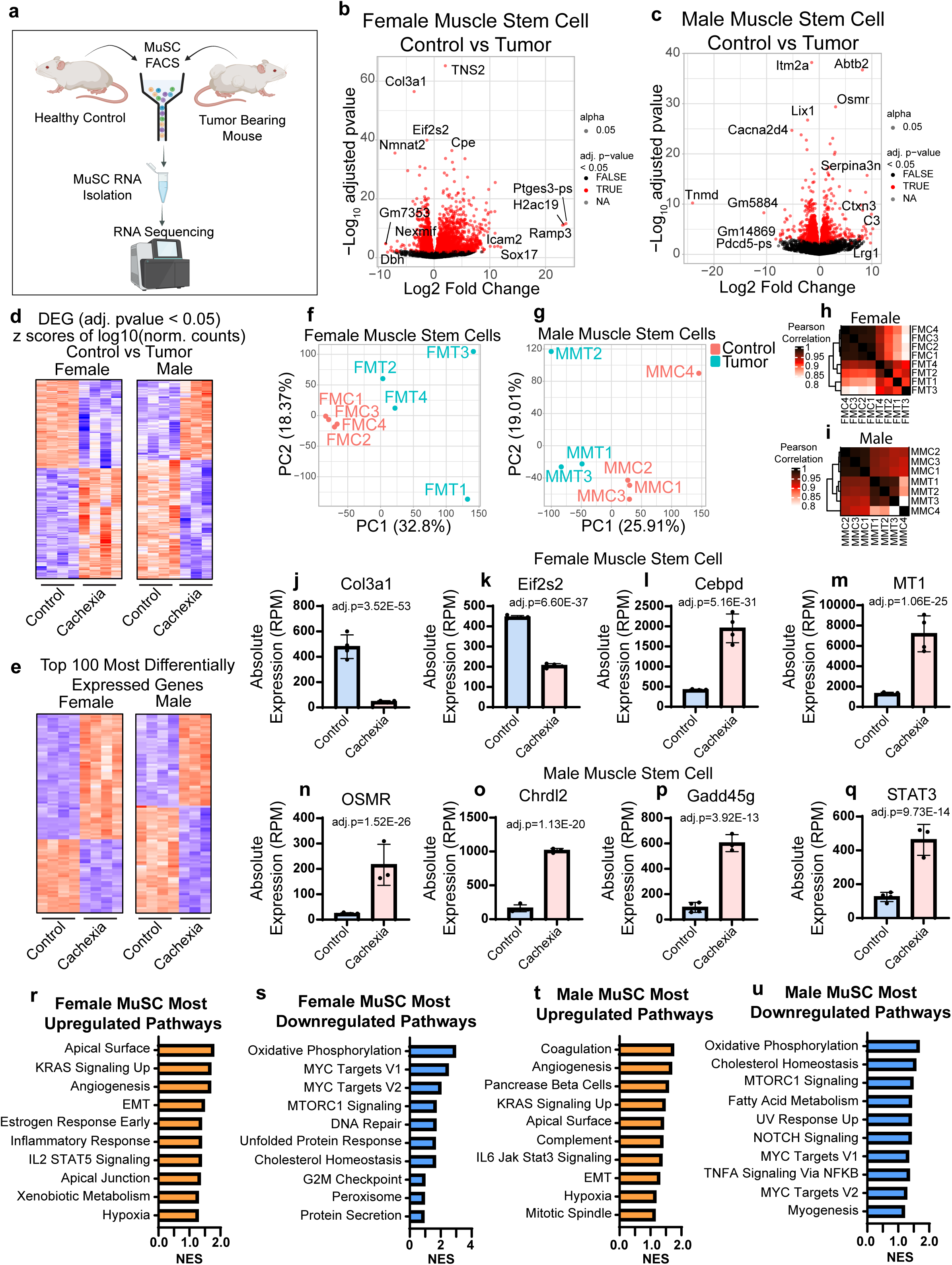
MuSCs are transcriptionally deregulated in cachexia. **a.** Schematic of the experimental design where pure populations of muscle stem cells are isolated from healthy controls and tumor bearing mice. The RNA from these cells are used to generate sequencing libraries. **b.** Volcano plot of the female MuSCs control vs tumor transcripts. **c.** Volcano plot of the male MuSCs control vs tumor transcripts. **d.** Heatmaps of the differentially expressed genes (adj. pvalue ≤ 0.05) of control vs tumor MuSC samples from both female and male mice. **e.** Heatmaps of the top 100 differentially expressed genes (adj. pvalue ≤ 0.05) of control vs tumor MuSC samples from both female and male mice. **f.** PCA plot of female MuSCs from control and tumor bearing mice. **g.** PCA plot of male MuSCs from control and tumor bearing mice. **h.** Pearson correlations between the sequenced female MuSC samples. **i.** Pearson correlations between the sequence male MuSC samples. **j.** RPM expression of Col3a1 in female MuSCs. **k.** RPM expression of Eif2s2 in female MuSCs. **l.** RPM expression of Cebpd in female MuSCs. **m.** RPM expression of MT1 in female MuSCs. **n.** RPM expression of OSMR in male MuSCs. **o.** RPM expression of Chrdl2 in male MuSCs. **p.** RPM expression of Gadd45g in male MuSCs. **q.** RPM expression of STAT3 in male MuSCs. **r.** Bar graph of the top ten most upregulated Hallmark pathways, in terms of enrichment score, from control vs tumor female MuSCs. **s.** Bar graph of the top ten most downregulated Hallmark pathways, in terms of enrichment score, from control vs tumor female MuSCs. **t.** Bar graph of the top ten most upregulated Hallmark pathways, in terms of enrichment score, from control vs tumor male MuSCs. **u.** Bar graph of the top ten most downregulated Hallmark pathways, in terms of enrichment score, from control vs tumor female MuSCs.

When comparing male and female MuSCs, we see that there is a lot of transcriptomic changes between the sexes (Extended Data Fig. 3a,b). The samples mostly cluster based on sex in both PCA plots and with Pearson Correlation, regardless of whether the sample came from healthy or cachectic mice (Extended Data Fig. 3c-f). We see that the most differentially expressed genes are typically Y chromosome genes, as was seen when comparing male and female myofibers. We wanted to assess whether the effect of cachexia was similar between male and female MuSCs. Using scatterplots plotting the LFC of gene expressing between control and cachectic, we see a strong correlation between the sexes when looking at both all genes and just the significantly different genes (Extended Data Fig. 3i,j)

Looking more in depth at the changes due to cancer in the MuSC transcriptome, we see in females that among the top 20 genes that are most significantly different, that Col3a1 is greatly reduced in cachexia (Fig. 3j). Col3a1 is an essential ECM component of the MuSC niche and its loss is associated with an increase in ECM stiffness^30^. Eif2s2 is also down in cachexia (Fig. 3k). This gene codes for a subunit of the eIF2 complex, and it is known that overexpression of this gene can induce muscle hypertrophy^31^, and the decrease in cachexia could therefore be contributing to muscle loss and a decrease in protein synthesis. Two other genes in the top 20 DEGs that are upregulated in cachexia are Cebpd and MT1 (Fig. 3l,m). Cebpd is a transcription factor that has a function in inducing inflammatory gene expression^32^. This may be one of the causes of the increase in inflammatory cytokine expression in MuSCs in cachexia, including Ccl11 and Il6 (data not shown). In male MuSCs, among the top 20 most differentially expressed genes, there are many genes important for proper MuSC function. We see that there is a large increase in OSMR expression (Fig. 3n). OSMR has multiple roles in MuSCs, particularly it is known to promote MuSC quiescence^33^, and it can also induce muscle atrophy^34^. Furthermore, Chrdl2 and Gadd45g are both increased in cachexia (Fig. 3o,p). Both of these genes have roles in maintaining MuSC quiescence. Chrdl2 is a known marker of quiescence^35,36^ and Gadd45g is a stress growth arrest factor^37^. We also see an increase in STAT3 expression in cachexia. STAT3 is a crucial transcription factor that has a plethora of functions in MuSCs, including self-renewal, proliferation and differentiation^38-40^. Inhibition of STAT3 has been shown to improve muscle regeneration in aging^41^. More broadly, when performing a GSEA analysis, we see that there is a lot of overlap between male and female MuSCs in the enrichment of key pathways. In both sexes, we see an upregulation of Kras Signaling and, importantly, the Epithelial Mesenchymal Transition pathway when comparing control vs cachexia MuSCs. Among the downregulated pathways, we see that Oxidative Phosphorylation and MTORC1 are enriched in both male and females (Fig. 3r-u).

We found that many of the genes that were altered due to cachexia, were also genes that were deregulated in aging and we wanted to determine whether there is a correlation between cachexia and aging. To confirm this, we compared the changes in the transcriptome of Control vs. Cachexia Male MuSCs to the changes in the transcriptome of Young vs. Aged Male MuSCs. The Young vs Aged data were retrieved from publicly available data (GSE171997). We see that there is a correlation between the effect of cachexia vs the effect of aging (Extended Data Fig. 4a,b). Specifically, we highlight here that both cachexia and aging cause a decrease in the expression of *Sparc* and *Col3a1*(Extended Data Fig. 4c,d), important genes for the assembly of the ECM of the MuSC niche. There is also a decrease in CCND1 (Extended Data Fig. 4e), which may indicate that the cachectic MuSCs have a delay in the entry into cycle. The inflammatory cytokine genes, *Ccl7* and *Ccl11*, are both highly upregulated in cachexia and aging (Extended Data Fig. 4f,g), which may be drivers of the increase in inflammation seen in both conditions. Lastly, Rnase4 is upregulated (Extended Data Fig. 4h), a gene that is involved in mRNA processing and could indicate changes to translation. Together, the data presented here demonstrates that cachexia induces large transcriptomic changes whose effects are multifaceted, affecting the ECM, inflammation, cell cycle and metabolism, among others.

### Cachexia Alters Chromatin Accessibility in MuSCs

Cachexia greatly affected the transcriptome of the MuSCs and we wanted to determine whether chromatin accessibility was also affected, as that can act as a regulator of gene expression. Using freshly isolated MuSCs from 4-month-old healthy control or tumor bearing female mice, we performed an ATAC-Seq (Fig. 4a). We see that cachexia greatly alters the chromatin accessibility of MuSCs. At a threshold of adjusted p-value ≤ 0.05, there are 6715 significantly differentially accessible peaks (Fig. 4b). Additionally, we do see that on a PCA plot the control and cachectic MuSCs cluster separately (Fig. 4c). We associated the peaks with the nearest gene and performed a GSEA analysis. We see that many of the Hallmark pathways that are enriched in the ATAC-Seq samples, are the same as the ones enriched in the RNA-Seq samples (Fig. 4d). For example, Kras Signaling, Inflammatory Response, Il6-Jak-Stat3 Signaling and the EMT pathways are all upregulated, as is the case with the RNA-Seq results. This would suggest that many of the transcriptomic changes seen in the MuSCs could be explained by alterations to chromatin accessibility and the chromatin state. Further supporting the effect of alterations in chromatin accessibility, we see that genes such as Myogenin and CCND1 have a decrease in accessibility in cachexia (Fig. 4e-g), while simultaneously having a decrease in mRNA expression. Both of these genes are important for MuSC function, with myogenin being necessary for differentiation and fusion into myofibers, while CCND1 promotes MuSC proliferation and expansion.

**Figure 4:**
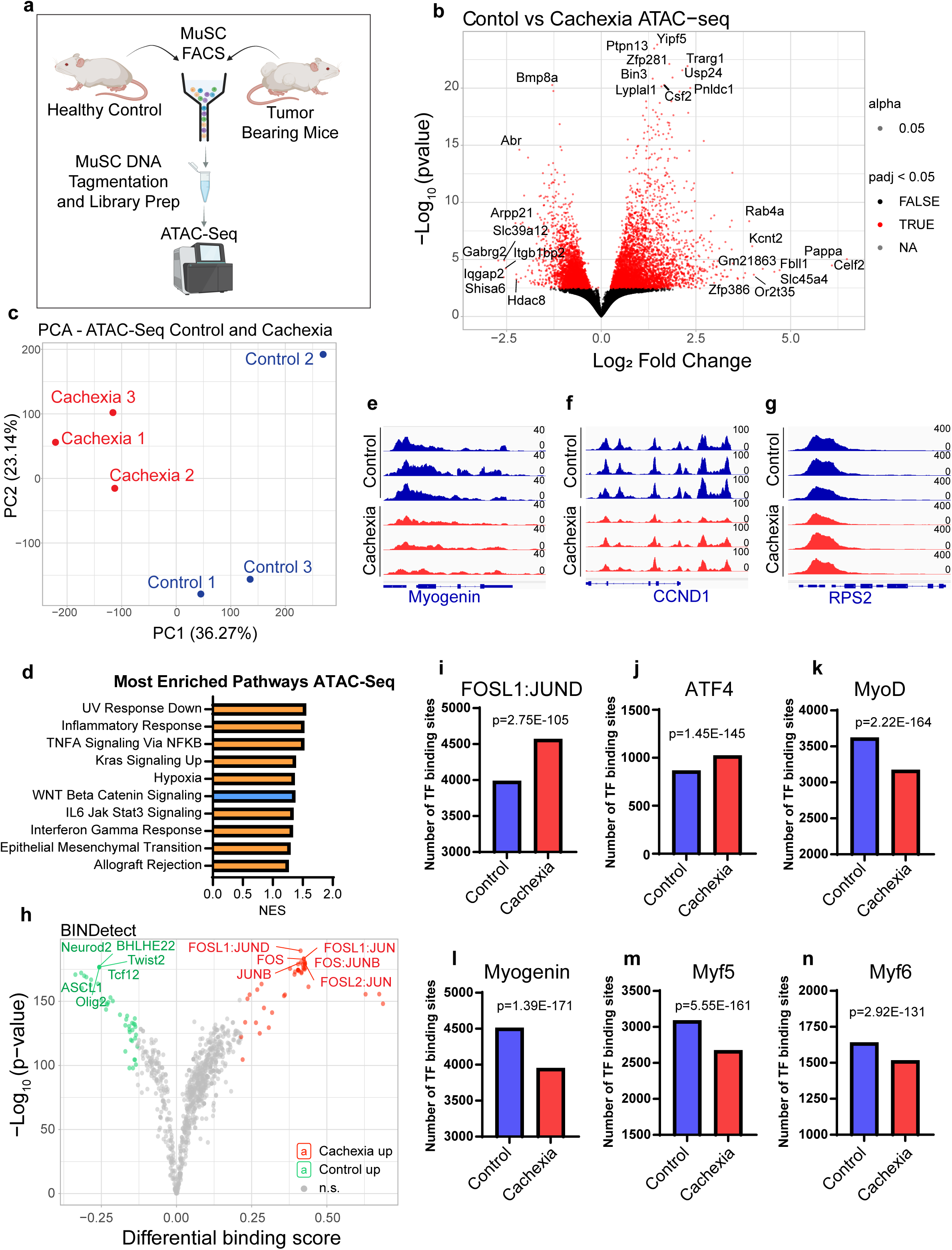
Epigenetic landscape is greatly altered in MuSCs from tumor bearing mice. **a.** Schematic of the experimental design where MuSCs from healthy and tumor bearing mice are FACS isolated. The DNA is tagmented and ATAC-Seq libraries are created from the isolated MuSCs. **b.** Volcano plot of the mapped peaks between MuSCs from healthy controls and cachectic mice. **c.** PCA plot of the control and cachectic MuSCs. **d.** The top ten most enriched pathways from the differentially accessible peaks between control and cachectic MuSCs. IGV tracks of ATAC-seq peaks at the **e**. Myogenin, **f**. CCND1, and **g**. RPS2 genes. **h.** Volcano plot of the changes in transcription factor binding in the differential peaks between control and cachectic MuSCs. Bar graphs of the number of transcription factor binding sites in control and cachectic MuSC peaks for **i.** FOSL:JUND, **j.** ATF4, **k.** MyoD, **l.** Myogenin, **m.** Myf5, and **n.** Myf6.

Lastly, we wanted to determine whether these changes in chromatin accessibility also led to changes in transcription factor binding. By analysing the DNA sequences under the differentially accessible peaks between control and cachexia MuSCs, we discovered that 88 transcription factors had a significant change in the number of binding motifs present in the peaks (Fig. 4 h). Many of the transcription factors that show an increase in their number of motifs are the Jun and Fos transcription factor families (Fig. 4h,i). The upregulation of these transcription factors is associated with stress response and inflammation has been determined to upregulate Fos expression^42^. Importantly, ATF4 has more accessible binding sites in cachexia than in healthy controls (Fig. 4j). Interestingly, we see that every myogenic regulatory factor (MRF) has a decrease in the number of accessible binding sites in the cachexia condition (Fig. 4k-n). Myf5, MyoD, Myogenin and Myf6 are all essential for the myogenic program to properly unfold. The loss of binding sites for the myogenic regulatory factors could be a major cause for MuSC dysfunction in cachexia.

### MuSCs Exhibit Functional Decline in Cachexia

To further elucidate whether MuSCs are functionally affected by cachexia, we isolated EDL myofibers and TA muscles from 4-month-old mice 2-3 weeks after C26 tumor cell injections. We found that EDL myofibers from cachectic mice had a significantly reduced number of MuSCs, and similarly, from the TA cross sections we see that there are also less MuSCs in cachexia compared to healthy controls (Fig. 5a-d). Importantly, it appears that there is a large decrease in the expression of PAX7 in MuSCs from tumor bearing mice. PAX7 is essential for the maintenance of the MuSC pool and a large decrease in its expression could be the primary cause of the loss in MuSC number and function^43^. However, it is important to note that this decrease in PAX7 protein is not reflected in the transcriptome. From the RNA-Seq, there is no significant decrease in *Pax7* expression. Therefore, it is probable that this decrease in PAX7 protein is due to translational changes in cachexia.

**Figure 5:**
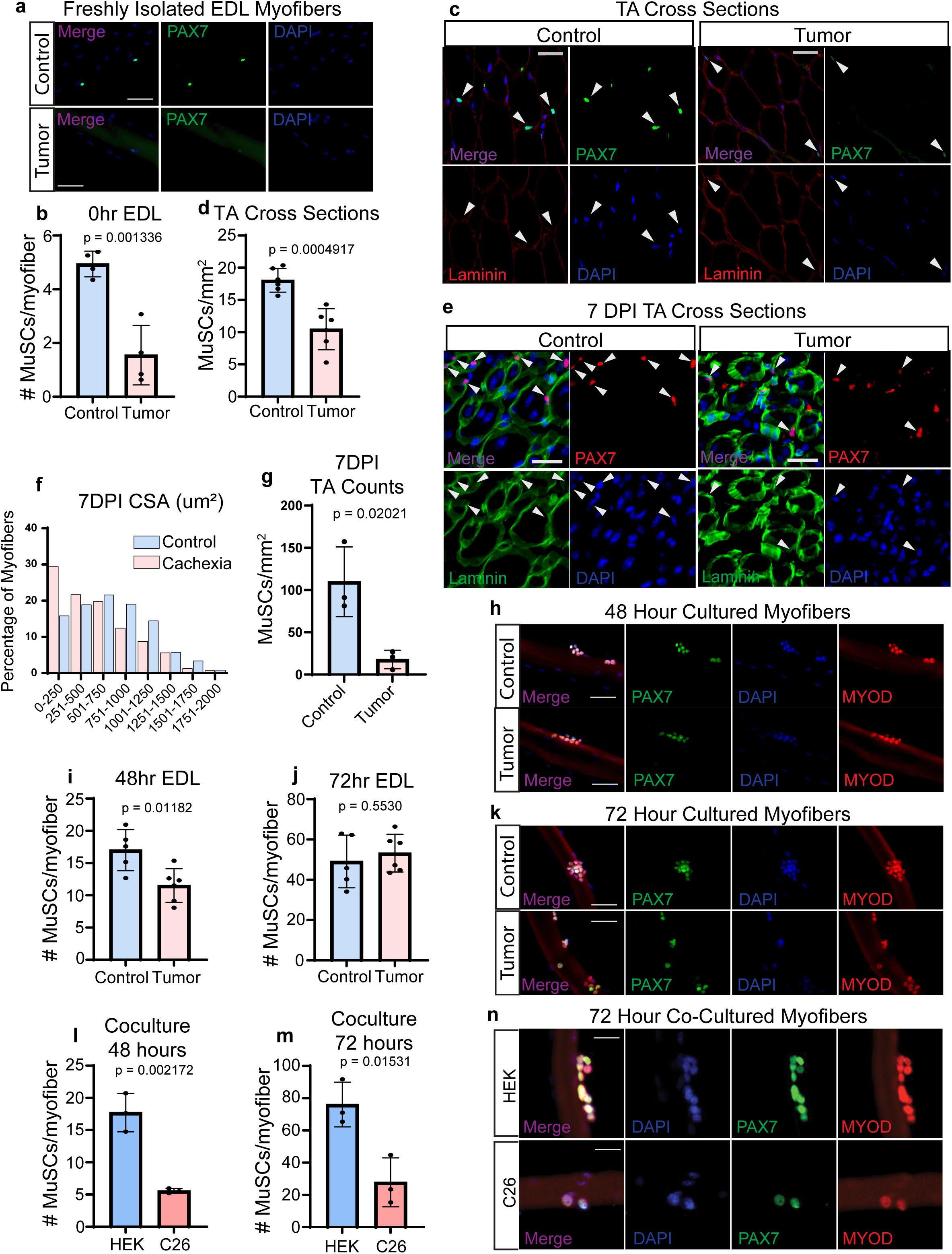
MuSCs from tumor bearing mice show a functional decline. **a.** Immunofluorescence staining of freshly isolated myofibers from control and tumor bearing female mice for PAX7. Scale bar = 30µm, n = 4 mice. **b.** Bar graph of the average number of MuSCs on freshly isolated myofibers from control and tumor bearing female mice. **c.** Immunofluorescence staining for PAX7 and Laminin on TA cross sections from control and tumor bearing female mice. Scale bar = 30 µm, n = 6 mice. **d.** Bar graph of the average number of MuSCs/mm2 in TA cross sections from control and tumor bearing female mice. **e.** Representative images of TA cross sections 7 days post injury (7DPI) from healthy control and tumor bearing female mice. Sections were stained for PAX7 and Laminin. Scale bar = 30 µm. n = 3 mice**. f.** CSA measurements of TA myofibers in control and tumor bearing female mice. **g.** Quantification of the number of MuSCs/mm2 in TA cross sections**. h.** Immunofluorescence staining of 48-hour cultured myofibers, from control and tumor bearing female mice, for PAX7 and MyoD. Scale bar = 30 µm, n = 5 mice. **i.** Bar graph of the average number of MuSCs on 48-hour cultured myofibers from control and tumor bearing female mice. **j.** Immunofluorescence staining of 72-hour cultured myofibers, from control and tumor bearing female mice, for Myogenin and MyoD. Scale bar = 30 µm, n = 5 control mice, n = 6 tumor bearing mice. **k.** Bar graph of the average number of MuSCs on 72-hour cultured myofibers from control and tumor bearing female mice. **l.** Bar graph of the average number of MuSCs on myofibers that were cocultured for 48 hours with either HEK cells or C26 cancer cells, n = 3 mice. **m.** Bar graph of the average number of MuSCs on myofibers that were cocultured for 72 hours with either HEK cells or C26 cancer cells. **n.** Representative immunofluorescence images of myofibers cocultured with HEK or C26 cancer cells for 72 hours. Myofibers were stained for PAX7 and MyoD. Scale bar = 20 µm.

To determine whether skeletal muscle regeneration was affected by C26 cancer, we injected 3-month-old mice with C26 cells. After 1 week of incubation, the TA of the mice were injured with cardiotoxin (CTX) and allowed to recover for 7 days. We see that the TAs of mice that had cancer did not regenerate as well as those from healthy controls (Fig. 5e). The regenerating fibers from cachexia are smaller and we see a large decrease in the number of MuSCs (Fig. 5f,g), which would further compromise the regenerative capacity of the muscle.

When MuSCs are removed from the tumor environment, they quickly recover. When we cultured isolated myofibers for 48 hours, we found that the MuSCs were capable of expanding and nearly reach the levels of the controls (Fig. 5h,i). By 72 hours, the number of MuSCs in the cachexia condition was equal to those in the control (Fig. 5j,k). These results indicate that MuSC function is altered due to cancer driven changes in the niche environment.

To determine whether C26 cells directly affect MuSC function, we performed a trans-well assay where isolated EDL myofibers were cocultured with C26 cells. Being exposed to C26 cells greatly affected the ability of the MuSCs to activate and proliferate. At 48 hours of coculture, there was almost no increase in the number of MuSCs on the EDLs cultured with C26 cells (Fig. 5l). This trend continued after 72 hours of coculture with C26 cells, with a significant decrease in the number of MuSCs (Fig. 5m,n). The same results were seen when the myofibers were cultured in C26 conditioned media. After 72 hours, the number of MuSCs on the myofibers in the conditioned media was significantly less than the number in the normal growth media, with numbers similar to those cultured in unsupplemented growth media (Extended Data Fig. 5a,b). Interestingly, we also saw a significant decrease in the number of Myogenin positive MuSC in the conditioned media (Extended Data Fig. 5a,c). To see whether this could be replicated in primary myoblasts, we cultured primary myoblasts in C26 conditioned media for 12, 24, and 36 hours. At all time points we found that there was a significant decrease in the percentage of Myogenin expressing cells (Extended Data Fig. 5d-i). Myogenin is the MRF that is responsible for the fusion of myoblasts into myotubes^44^, as such we wanted to determine whether this observed decrease in Myogenin would translate into deficient differentiation. We cultured primary myoblasts for 5 days in C26 conditioned differentiation media to assess the effect on Myogenin expression and the fusion of these cells. We found that the conditioned media resulted in a decrease in Myogenin positive nuclei, consistent with our findings in the primary myoblasts (Extended Data Fig. 5j,k). We also noted a decrease in the fusion of the cells into myotubes and a decrease in the size of the myotubes themselves (Extended Data Fig. 5l-m). All together, the data presented here indicates that C26 cancer cells are capable of directly communicating with MuSCs. This communication has a broad deleterious effect on the myogenic program, reducing expansion of the MuSC pool and impeding differentiation.

### C26 Tumor Cells Adapt to the Host Environment

To further elucidate the communication network between C26 cells and MuSCs, we needed to assess the transcriptome of the C26 cells. We also wanted to determine whether the transcriptome of the C26 cells was altered when the cells were engrafted and grown *in vivo* compared to cultured *in vitro*, as well as whether the sex of the host mouse had any bearing. To achieve this goal, we created a GFP over expressing C26 cell line through retroviral infection. The GFP cells were injected into 3-month-old male and female Balb/c mice and allowed to form tumors (Fig. 6a). The tumors were grown until the mice neared endpoint. At that time the tumors were dissected and single cell suspensions created in order to FACS sort the GFP positive cells (Fig. 6b,c). RNA-Seq libraries were generated from the cultured *in vitro* GFP C26 cells, as well as the GFP C26 cells isolated from male and female mice.

**Figure 6:**
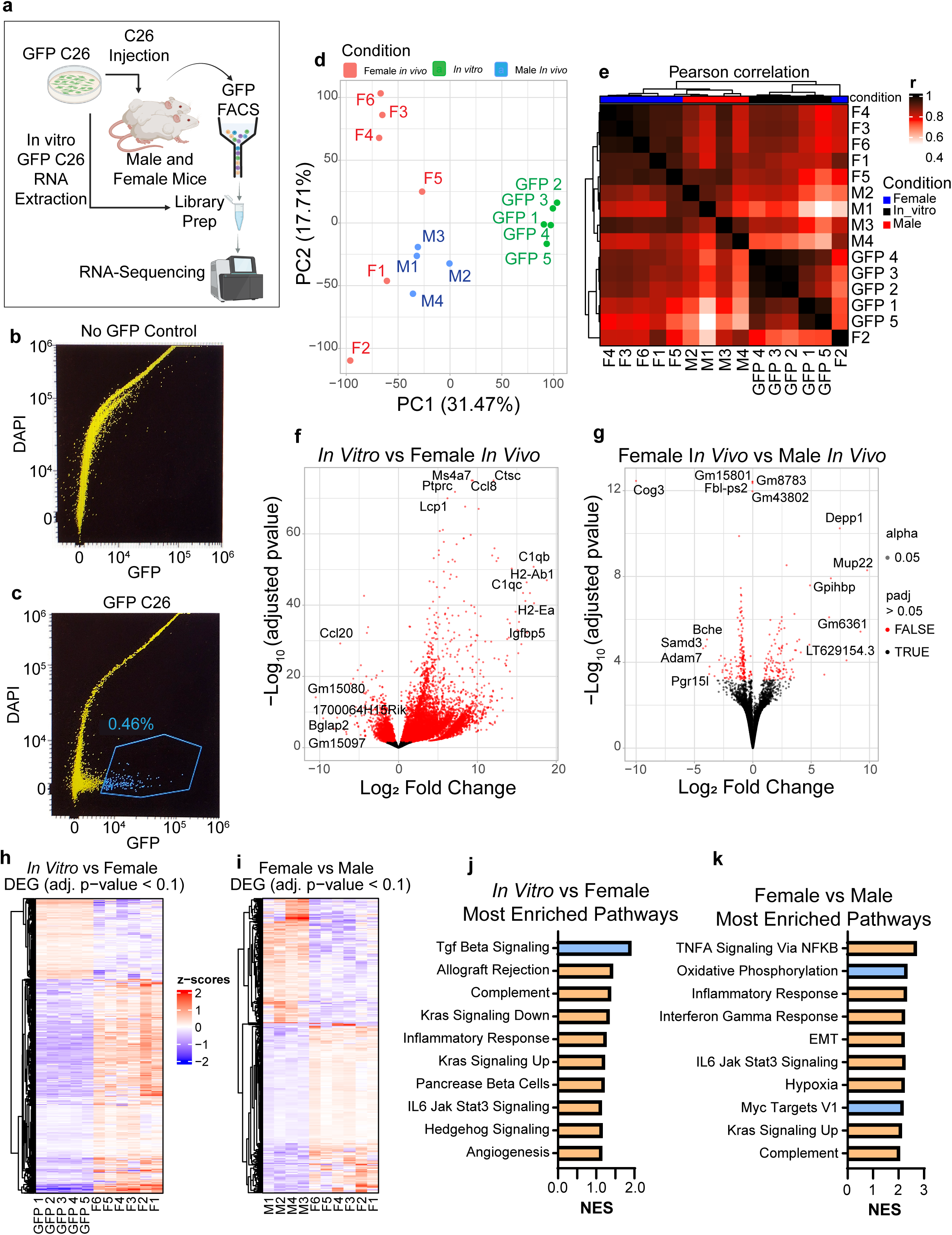
C26 cancers cells alter their transcriptome in an *in vivo* environment. **a.** Schematic of the experimental design, where C26 cells are transformed to stably express GFP. The cells are injected into both male and female Balb/c mice. The GFP cells are reisolated from the solid tumors and, simultaneously with the *in vitro* cultured cells, RNA-Sequencing libraries are generated. Figure created with Biorender.com. **b.** FACS plot of No-GFP control, indicating GFP and DAPI intensity. **c.** FACS plot of GFP C26 solid tumors and gating for the isolation of GFP positive C26 cells. **d.** PCA plot of RNA-Seq C26 samples from *in vitro* culture, *in vivo* in female mice and *in vivo* in male mice. **e.** Pearson correlation graph between the *in vitro* and *in vivo* samples. **f.** Volcano plot of the difference in transcript expression between *in vitro* cultured C26 cells and female *in vivo* C26 cells. **g.** Volcano plot of the difference in transcript expression between female *in vivo* and male *in vivo* C26 cells. **h.** Heatmap of differentially expressed genes (adj. pvalue ≤ 0.1) between *in vitro* and female *in vivo* C26 cells. **i.** Heatmap of differentially expressed genes (adj. pvalue ≤ 0.1) between female *in vivo* and male *in vivo* C26 cells. **j.** Bar graph of the top ten most enriched Hallmark pathways between *in vitro* C26 and *in vivo* C26 in female mice. Blue bars represent downregulated pathways; orange bars represent upregulated pathways. **k.** Bar graph of the top ten most enriched Hallmark pathways between *in vivo* C26 in female mice and *in vivo* C26 in male mice. Blue bars represent downregulated pathways; orange bars represent upregulated pathways.

From the sequencing results, we see that there is a large change in the transcriptome when the C26 cells were grown *in vivo*, with the *in vivo* samples clustering away from the *in vitro* ones (Fig. 6d,e). Interestingly, there are far more genes that are differentially expressed between the *in vitro* and *in vivo* samples than when comparing male and female grown cells. At a threshold of adjusted p-value ≤ 0.1, there are 7880 upregulated and 3155 downregulated genes between *in vitro* and *in vivo* in females, while only having 200 upregulated and 272 downregulated genes between female and male *in vivo* C26 cells (Extended Data Fig. 6a). The results are even more striking when using our most stringent threshold of adjusted p-value ≤ 0.001. Between *in vitro* and *in vivo* in females there are 4708 and 907 upregulated and downregulated genes, respectively. However, between *in vivo* female and male, there is a paltry 11 genes that are significantly upregulated and 20 that are significantly downregulated (Extended Data Fig. 6a). The sex of the host appeared to have a minimal effect on the transcriptome of the C26 cells, when compared to the effect seen between *in vitro* and *in vivo* (Fig. 6f-i; Extended Data Fig. 6b,c). From the GSEA analysis, we see that there is a large overlap in the enriched Hallmark pathways when comparing *in vitro* vs *in vivo* in female and *in vitro* vs *in vivo* in male. Among the top enriched pathways, Allograft Rejection, Complement, Kras Signaling, and Il6 Jak Stat3 Signaling are upregulated in both comparisons (Fig. 6j; Extended Data Fig. 6d). We also see that Inflammatory Response, EMT and Il6 Jak Stat3 Signaling pathways are enriched when comparing *in vivo* in female vs *in vivo* in male (Fig. 6k). Together, these data indicate that C26 cells adapt to the *in vivo* environment to promote their survival and growth, but that the sex of the host has no major effect on C26 behaviour.

Our results indicate that many immune cell genes are upregulated in the *in vivo* condition compared to the *in vitro*. To ensure that the changes in the transcriptome previously observed were not due simply to host cell contamination or our RNA-Seq, we further assessed the expression of immune related genes. We found that while there was an upregulation of many immune genes *in vivo*, including *Adgre1* and *Mrc1*, many immune genes were not expressed at all, such as *CD163* and *MARCO* (Extended Data Fig. 6e). This makes it unlikely that there is immune cell contamination as there would not be selective expression of certain immune related genes, but rather an increase in all the immune genes. Interestingly, some immune genes were already expressed in the *in vitro* cultured condition, in which there could not be any immune cell contamination, including CD14 and CD80 (Extended Data Fig. 6e). To further confirm there are no host contaminating cells in the samples, we analyzed the expression of Y chromosome genes. We chose to do this because C26 cells were originally derived from female Balb/c mice and are therefore female cells lacking a Y chromosome. If there were to be contamination from the host, we would see expression of Y chromosome genes in the *in vivo* in male mice condition. However, that is not the case, there is no upregulation of Y chromosome genes in the *in vivo* in male condition compared to *in vitro* and *in vivo* in female (Extended Data Fig. 6f,g). Using our male and female MuSC RNA-Seq data for comparison, we see that only the male MuSCs express Y chromosome genes, with the C26 cells resembling the female MuSCs (Extended Data Fig. 6f). These results indicate that there is no contamination from other cell types, and that it is the C26 cells that are expressing these immune genes. We speculate that the C26 cells are undergoing immune mimicry to evade the host immune system.

### Cancer Induces an Increase in Circulating MMP9 and GDF15 Which Impedes MuSC Function

C26 tumors are solid tumors that we have injected subcutaneously and therefore are not present in the muscle itself. Any communication that C26 cells have with muscle cells must therefore be through the production of signaling molecules that travel through the blood. To assess how the cytokine landscape is altered in cachexia, we performed a Cytokine Array on serum from male and female mice that were either healthy or injected with C26 tumors. Several cytokines changed their levels due to cachexia, with a particular interest in MMP9 and GDF15 (Fig. 7a,b). From the array, we see that in cachexia MMP9 levels are approximately 5 times and 3 times higher in male and female mice, respectively (Extended Data Fig. 7a,b). Similarly, GDF15 is approximately 25 times higher in cachectic male mice compared to controls, and approximately 4 times higher in cachectic females (Fig. 7c,d). These cytokines are produced by the C26 cells, with MMP9 being expressed only in the C26 cells, not in MuSCs or Myofibers, and increasing in transcriptional expression when C26 is grown *in vivo* (Extended Data Fig. 7c). GDF15 is produced in both the C26 cells and MuSCs, with its expression transcriptionally decreasing when C26 cells are grown *in vivo* (Fig. 7e,f).

**Figure 7:**
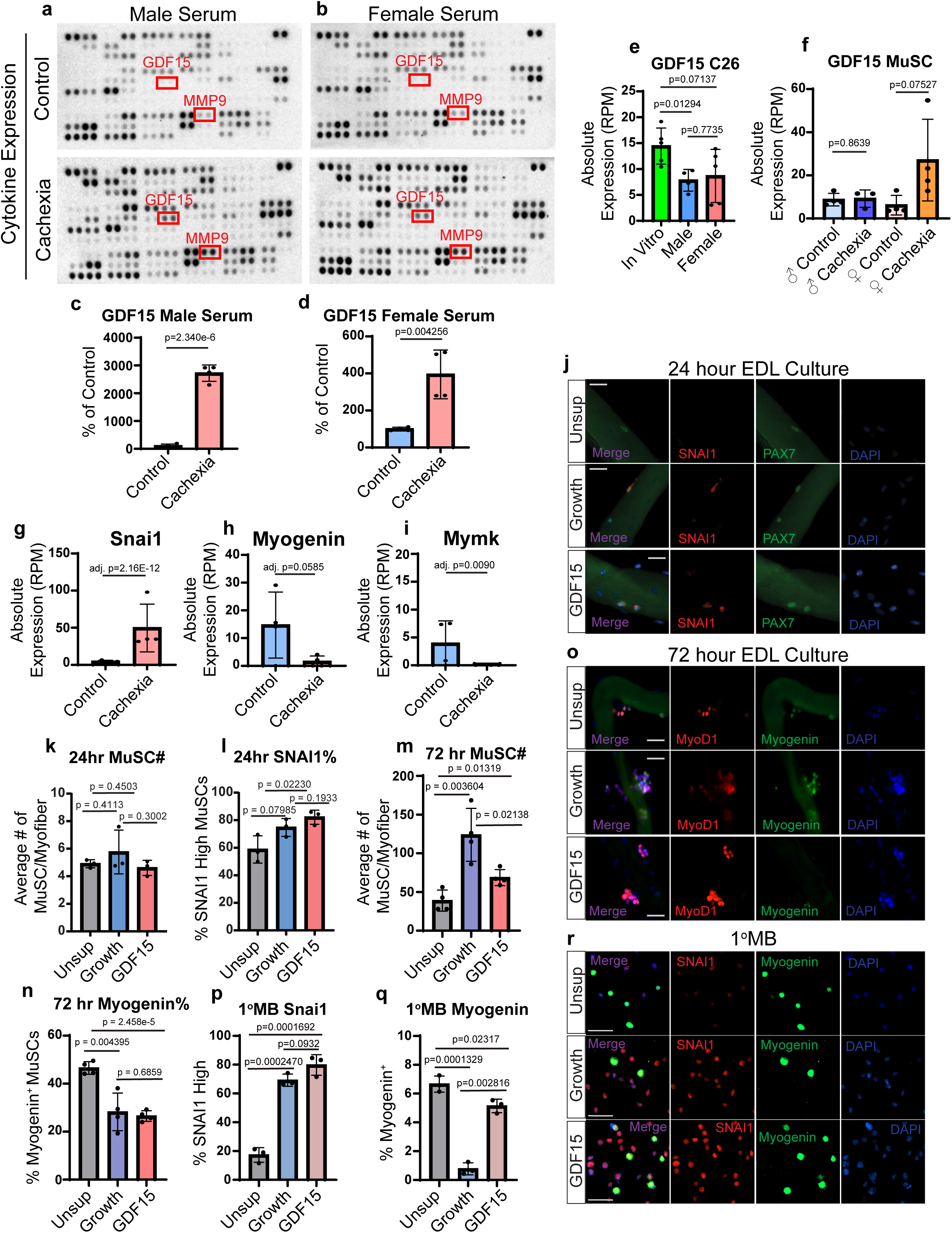
GDF15 induces SNAI1 in muscle stem cells and blocks differentiation. **a.** Membranes from Proteome Profiler Mouse XL Cytokine Array incubated with serum collected from 4-month-old male healthy control balb/c mice and tumor bearing mice 14 days post C26 injection, n = 2 mice. **b.** Membranes from Proteome Profiler Mouse XL Cytokine Array incubated with serum collected from 4-month-old female healthy control balb/c mice and tumor bearing mice 14 days post C26 injection, n = 2 mice. **c.** Quantification of GDF15 level in the serum from male healthy control and tumor bearing mice. Values are normalized and presented as a percentage of the control. **d.** Quantification of GDF15 level in the serum from female healthy control and tumor bearing mice. Values are normalized and presented as a percentage of the control. **e.** Bar graph of the absolute expression of GDF15 transcript (RPM) in GFP-C26 cells cultured *in vitro* or isolated *in vivo* from male and female tumor bearing mice. **f.** Bar graph of the absolute expression of GDF15 transcript (RPM) in MuSCs freshly isolated from male and female 3-month-old healthy control and tumor bearing mice. Bar graph of the absolute expression of **g.** Snai1, **h.** Myogenin, and **i.** Mymk transcripts (RPM) in MuSCs freshly isolated from female 3-month-old healthy control and tumor bearing mice. **j.** Representative pictures of isolated EDL myofibers cultured for 24 hours in unsupplemented growth media (15% FBS, 1% P/S, – CEE, – bFGF), normal growth media (15% FBS, 1% P/S, 1% CEE, 5 ng/µL bFGF) and GDF15 supplemented growth media (15% FBS, 1% P/S, 500 ng/µL GDF15, – CEE, – bFGF), GDF15 was added every 12 hours. Myofibers were stained for PAX7 and SNAI1. n = 4 mice. Scale bar = 20 µm. **k.** Quantification of the average number of MuSCs per EDL myofiber from myofibers cultured for 24 hours in unsupplemented, normal growth and GDF15 treated conditions. **l.** Quantification of the percentage of high SNAI1 expressing MuSCs per EDL myofiber from myofibers cultured for 24 hours in unsupplemented, normal growth and GDF15 treated conditions. **m.** Quantification of the average number of MuSCs per EDL myofiber from myofibers cultured for 72 hours in unsupplemented, normal growth and GDF15 treated conditions. **n.** Quantification of the percentage of Myogenin expressing MuSCs per EDL myofiber from myofibers cultured for 72 hours in unsupplemented, normal growth and GDF15 treated conditions. **o.** Representative pictures of isolated EDL myofibers cultured for 72 hours in unsupplemented growth media (15% FBS, 1% P/S, – CEE, – bFGF), normal growth media (15% FBS, 1% P/S, 1% CEE, 5 ng/µL bFGF) and GDF15 supplemented growth media (15% FBS, 1% P/S, 500 ng/µL GDF15, – CEE, – bFGF), GDF15 was added every 12 hours. n = 4 mice. Scale bar = 20 µm. **p.** Quantification of the percentage of high SNAI1 expressing primary myoblasts that were cultured for 24 hours in unsupplemented, normal growth and GDF15 treated conditions. **q.** Quantification of the percentage of Myogenin expressing primary myoblasts that were cultured for 24 hours in unsupplemented, normal growth and GDF15 treated conditions. **r.** Representative pictures of primary myoblasts cultured for 24 hours in unsupplemented (20% FBS, 1% P/S), normal growth (20% FBS, 1% P/S, 5 ng/µL bFGF), or GDF15 treated (20% FBS, 1%P/S, 500 ng/µL GDF15) media. The GDF15 was added every 12 hours. Primary myoblasts were stained for SNAI1 and Myogenin. n = 3 biological replicates. Scale bar = 50 µm.

From the cancer genome atlas (TCGA) we retrieved human data from patients with colon adenocarcinoma and pancreatic adenocarcinoma. Only patients that did not receive chemotherapy or surgery were selected. From these data, we see in both types of cancer that the patients that had higher levels of GDF15 or MMP9 had a worse prognosis than those that had lower levels of the cytokines (Extended Data Fig. 7d-g). These findings highlight the potential role of GDF15 and MMP9 in cancer progression.

Both GDF15 and MMP9 are known to induce the expression of SNAI1 and upregulated the EMT pathway. This is of particular interest as we see from the RNA-Seq data that the EMT pathway is upregulated in cachectic MuSCs, and specifically we see a large upregulation of *Snai1* expression (Fig. 7g). Previous studies have shown that SNAI1 represses myogenic differentiation by outcompeting MyoD binding and blocking the expression of Myogenin^45^. An aberrant expression of SNAI1 can block the ability of myogenic cells from differentiating and fusing into myotubes. These findings are corroborated by our MuSC RNA-Seq data, where we see a decrease in Myogenin expression and key differentiation genes, such as *Mymk*, which is necessary for myoblast fusion (Fig. 7h,i).

To determine whether MMP9 can directly induce SNAI1 expression in primary myoblasts, we treated these cells with MMP9 for 12 hours. Staining revealed that there was a higher percentage of High SNAI1 expressing cells in the MMP9 treated condition compared to the controls (Extended Data Fig. 8a,b). However, we did not observe a decrease in Myogenin at this timepoint (Extended Data Fig. 8c). Further, we treated EDL myofibers, that were cultured 24 hours prior, with MMP9 for 1 hour and then stained for phosphor-SMAD3 (Extended Data Fig. 8d). Exposure to MMP9 resulted in an increase in phosphor-SMAD3 staining compared to controls (Extended Data Fig. 8e). These results suggest that MMP9 can directly signal to myogenic cells and induce the expression of SNAI1.

We also wanted to determine whether GDF15 can directly signal to MuSCs. Freshly isolated EDL myofibers were cultured in unsupplemented growth media and treated with GDF15. 24 hours post isolation, we see that while there was no change in the MuSC number, there is an increase in the percentage of High SNAI1 expressing MuSCs (Fig. 7j-l). However, at 60 and 72 hours of culture, we see that compared to the unsupplemented myofibers, the GDF15 treatment resulted in an increase in MuSC number and a decrease in Myogenin expressing MuSCs (Fig. 7m-o; Extended Data Fig. 8f-h). Primary myoblasts were then treated for 24 hours with GDF15, and similar results were obtained. Primary myoblasts that were treated with GDF15 had higher levels of SNA1 and lower levels of Myogenin compared to the unsupplemented condition (Fig. 7p-r). Together, these data indicate that GDF15 can affect MuSC function, promoting proliferation at the expense of differentiation.

### Knockout of GDF15 in C26 Cells does not Prevent Cachexia

GDF15 has recently become a protein of great interest in the field of cachexia research. Recent studies have shown that GDF15 can suppress the appetite of cancer patients and that, by blocking GDF15 in the body, cachexia can be prevented^46^. We wanted to determine whether knocking out GDF15 in C26 tumors can prevent muscle wasting in mice. To achieve this, we generated Crispr GDF15 knockout and Crispr Control C26 cells. Mice were subcutaneously injected with the Crispr cells and tumors were allowed to grow until mice reached endpoint. Surprisingly, the GDF15 KO tumors grew faster, and mice reached endpoint within 3 weeks, compared to Crispr Control tumors where mice who reached endpoint between 4-5 weeks post tumor injection. However, this does not appear to be due to differences in proliferation between the cells as *in vitro* EDU assays showed equal EDU incorporation (Extended Data Fig. 9a,b) The KO of GDF15 in the C26 cells had no effect on body mass, with both the KO and the controls losing the same amount of body weight (Fig. 8a). The GDF15 KO tumors grew larger than the controls (Fig. 8b) and interestingly, the TA muscle mass was significantly reduced in the GDF15 KO tumor condition compared to the Crispr Control mice (Fig. 8c). The TA cross sectional area of the GDF15 KO tumor mice was slightly reduced compared to the Crispr controls as well (Fig. 8d,e). Overall, there was no significant difference between the percentage of fat mass and lean mass between the Crispr Control and GDF15 KO tumor bearing mice (Fig. 8f,g). We also validated the levels of circulating GDF15 and found that while there was a significant reduction of GDF15 in the serum of the Crispr KO tumor bearing mice, there was still GDF15 present, meaning that the tumor is not the sole source of GDF15 (Fig. 8h). Furthermore, the KO of GDF15 in the tumor did not result in any differences in metabolism or food consumption during the second week post tumor injection (Fig. 8i,j). We do see, surprisingly, that there was a trend towards a lower daily food intake in the GDF15 KO tumor bearing mice (Fig. 8i). This is contrary to expectations as GDF15 is a known appetite suppressor. However, as previously stated, the GDF15 KO tumors grew faster and at this time point the mice in the GDF15 KO cohort were nearing endpoint, while the Crispr Controls were still relatively healthy. This could explain the discrepancy. We also observed a trend of the GDF15 KO tumor bearing mice shifting their energy consumption towards fat instead of carbohydrates, as evidenced by the decrease in RER towards 0.7.

**Figure 8:**
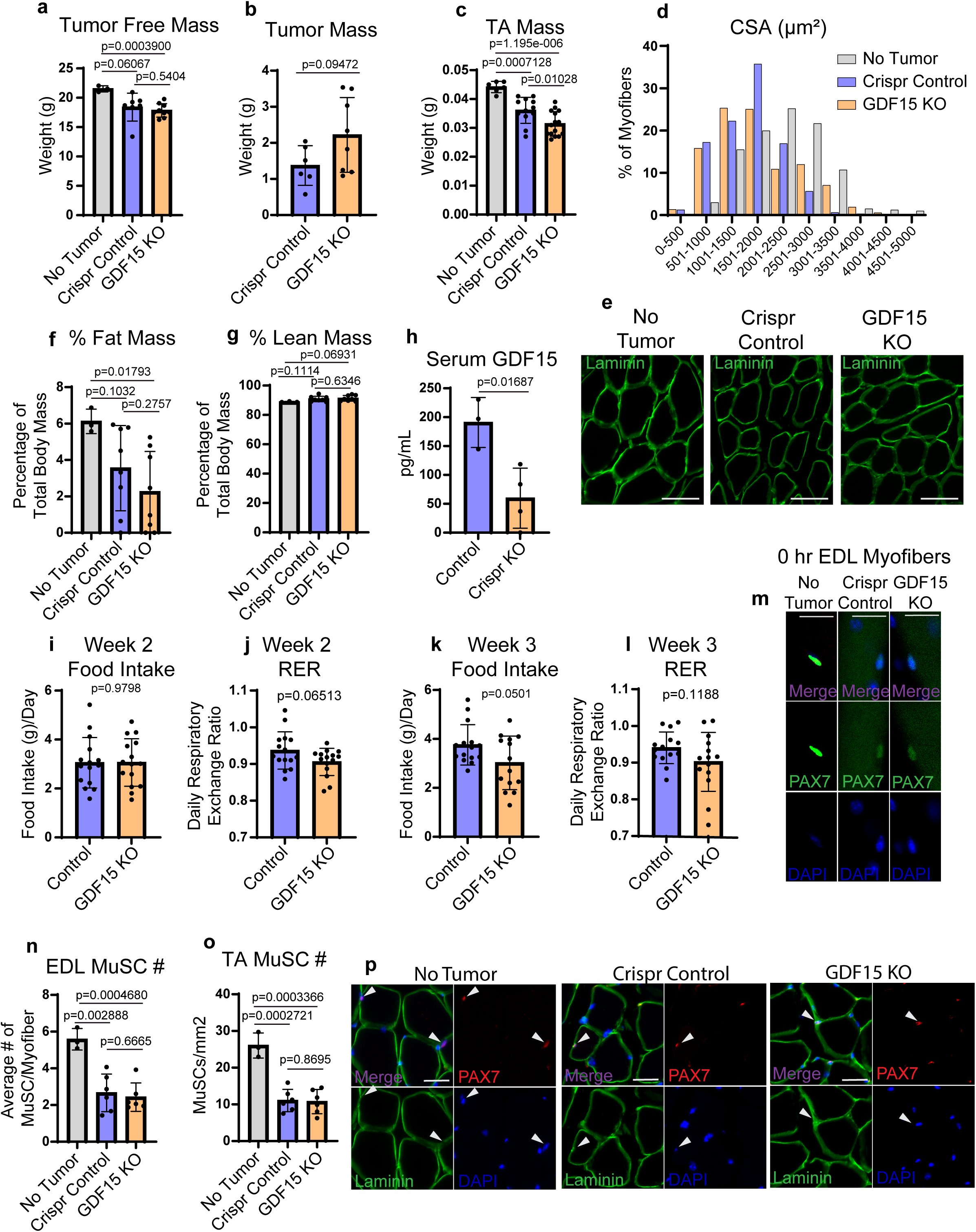
Tumor derived GDF15 is not a primary cause of muscle wasting or muscle stem cell dysfunction. **a.** Bar graph of the tumor free mass of female balb/c mice that are either healthy controls, mice injected with Crispr Control C26 cancer cells, or mice injected with Crispr GDF15 KO C26 cancer cells. All tumor bearing mice were injected with C26 cells at 3 months of age. Tumor free mass was assessed when mice reached endpoint, healthy controls were collected at the end of the experiment. n = 3 healthy controls, 6 Crispr Controls and 8 GDF15 KO. **b.** Bar graph of the mass of the tumors isolated from the Crispr Control and GDF15 KO tumor bearing mice. n = 6 Crispr Controls and 8 GDF15 KO. **c.** Bar graphs of the mass of the tibialis anterior (TA) isolated from healthy controls, Crispr Control and GDF15 KO tumor bearing mice. n = 6 TAs from healthy controls, 12 TAs from Crispr Controls, and 16 TAs from GDF15 KO. **d.** Quantification of the cross-sectional area (CSA) of myofibers in the TA isolated from Healthy Controls, Crispr Controls and GDF15 KO tumor bearing mice. Data is presented as percentage of myofibers that are within the marked size range. n = 3 healthy controls, 6 Crispr Controls and 6 GDF15 KO tumor bearing mice. **e.** Representative picture of the CSA of myofibers from the TA of Healthy Control, Crispr Control, and GDF15 KO tumor bearing mice. Cross sections were stained for Laminin. Scale bar = 80 µm. **f.** Quantification of fat mass as a percentage of whole body mass, measured by EchoMRI 19 days post tumor cell injection. **g.** Quantification of lean mass as a percentage of whole body mass, measured by EchoMRI 19 days post tumor cell injection. **h.** Quantification of the level of serum GDF15, measured by ELISA, from Crispr Control and GDF15 KO tumor bearing mice. Blood was collected 14 days post tumor injection. n = 4 mice. **i.** Daily food intake of Cripsr Control and GDF15 KO tumor bearing mice. Food intake was measured using a Comprehensive Lab Animal Monitoring System (CLAMS) for 3 days in the second week post tumor injection. n = 5 mice. **j.** Daily Respiratory Exchange Rate (RER) of Cripsr Control and GDF15 KO tumor bearing mice. RER was measured using a CLAMS for 3 days in the second week post tumor injection. n = 5 mice. **k.** Daily food intake of Cripsr Control and GDF15 KO tumor bearing mice. Food intake was measured using a Comprehensive Lab Animal Monitoring System (CLAMS) for 3 days in the third week post tumor injection. n = 5 mice. **l.** Daily Respiratory Exchange Rate (RER) of Cripsr Control and GDF15 KO tumor bearing mice. RER was measured using a CLAMS for 3 days in the third week post tumor injection. n = 5 mice. **m.** Representative immunofluorescence pictures of freshly isolated EDL myofibers from Healthy Controls, Crispr Control and GDF15 KO tumor bearing mice. Myofibers were stained for PAX7. n = 3 Healthy Controls, 6 Crispr Controls and 6 GDF15 KO tumor bearing mice. Scale bar = 20 µm. **n.** Quantification of the average number of muscle stem cells per myofiber from Healthy Controls, Crispr Control and GDF15 KO tumor bearing mice. n = 3 Healthy Controls, 6 Crispr Controls and 6 GDF15 KO tumor bearing mice. **o.** Quantification of the number of muscle stem cells per mm^2^ in TA cross sections from Healthy Controls, Crispr Controls and GDF15 KO tumor bearing mice. **p.** Representative pictures of immunofluorescence staining of TA cross sections from Healthy Controls, Crispr Controls and GDF15 KO tumor bearing mice. Cross sections were stained for Laminin and PAX7. n = 3 Healthy Controls, 6 Crispr Controls and 6 GDF15 KO tumor bearing mice. Scale bar = 30 µm.

Next, we assessed the effect of knocking out GDF15 in the C26 tumors on MuSC numbers. Freshly isolated EDL myofibers were stained for Pax7 and the MuSC number quantified. We found that there was no difference between the Crispr Control and GDF15 KO conditions, both exhibited a similar reduction in MuSC number when compared to healthy controls (Fig. 8m,n). We also quantified the number of MuSCs in TA cross sections, and the same result was obtained, both tumor conditions resulted in an equal loss of MuSCs (Fig. 8o,p). We then assessed how primary myoblasts cocultured with GDF15 KO C26 cells were affected. We found that there was no change in the proliferation of primary myoblasts, as evidenced by EDU incorporation (Extended Data Fig. 9c,d). Further, differentiation was equally reduced when cells were differentiated in the presence of Crispr Control or GDF15 KO C26 cells (Extended Data Fig. 9e,f). We did observe, however, an increase in the number of MuSCs in the EDL myofibers that were cocultured with the GDF15 KO C26 cells when compared to those cultured with the Crispr Control cells, as well as a reduction in Myogenin positive MuSCs (Extended Data Fig. 9g-i). As a whole the data presented here indicates that, in mice, the knockout of GDF15 in the tumor is not sufficient to rescue the cachexia phenotype and does not prevent the loss of MuSCs.

## DISCUSSION

Cancer cachexia leads to a dramatic alteration of skeletal muscle encompassing the myofiber to its resident stem cells. Through an integrative multi-pronged approach using single-myofiber and MuSC RNA-seq, ATAC-seq, proteomic tracing, and regeneration assays, we demonstrate that cachexia represents a global suppression of the myogenic differentiation program concurrent with induction of muscle atrophy. Our data suggest that suppression of muscle stem cell differentiation arises from altered transcriptional, metabolic, and epigenetic remodeling driven by tumor-derived secreted factors, such as GDF15 and MMP9.

Single-myofiber RNA-seq shows that cachexia drives extensive transcriptional alterations leading to significant changes in their genetic network related to metabolic state and stress response. The upregulation of PDK4, ACOT1, and ACOT2 suggests a shift toward fatty-acid oxidation, consistent with decreased respiratory exchange ratio and fat mass observed *in vivo* (Fig. 2). Additionally, downregulation of Mafa, Myh4, and Ampd1 suggest selective susceptibility of fast-twitch fibers to cachexic environment (Fig. 2). Enrichment of hypoxia and apoptotic pathways, coupled with repression of WNT and Notch signaling, highlights a reduction of anabolic programs within the myofibers. These alterations are in line with human cachexia studies implicating oxidative stress, mitochondrial dysfunction, and metabolic reprogramming as important drivers of atrophy^47-50^.

In the cachectic environment MuSCs exhibit reduced expression of extracellular-matrix and niche-organizing factors such as Col3a1 and Sparc, respectively and key regulators of proliferation and translation (CCND1, Eif2s2), alongside induction of inflammatory and quiescence-associated transcripts (Ccl7, Il6, Osmr, Chrdl2, Stat3) (Fig. 3). Pathway enrichment analysis implicates EMT, IL6–JAK–STAT3, and KRAS signaling, and suppression of oxidative phosphorylation and mTORC1 activity for muscle stem cells. These data collectively suggest a chronic stress-induced, regeneration deficient state. Interestingly, comparison of gene expression data with aged MuSCs reveals a substantially common transcriptomic signature between cachectic and aged muscle stem cells, suggesting that cachexia phenocopies aspects of muscle stem cell aging through activation of inflammatory pathways and metabolic reprogramming (Extended Data Fig. 4).

ATAC-seq provides further mechanistic insight into this reprogramming. In freshly sorted muscle stem cells, cachexia reduces chromatin accessibility at the Myogenin and CCND1 loci while enhancing the accessibility of ATF4, FOS and JUN binding motifs. These findings suggest that stress-responsive factors dominate the cachectic regulatory landscape, concurrent with the repression of the myogenic differentiation network. Such differentiation related chromatin erosion mirrors observations in aging and dystrophic contexts where environmental stress remodels stem-cell epigenomes^51-53^. Although MuSCs from cachectic mice display impaired proliferation and regenerative capacity, they rapidly regain their function *ex vivo*, underscoring that their dysfunction is largely reversible and niche-driven rather than due to cell-intrinsic dysregulation (Fig. 5).

Transcriptomic profiling of GFP-labeled C26 cells before and after injection into the mice revealed a dramatic *in vivo* reprogramming, including activation of IL6–JAK–STAT3, KRAS, and EMT pathways, as well as induction of Adgre1 and Mrc1. These changes were independent of host sex, suggesting that tumor adaptation to the host microenvironment rather than hormonal milieu drives cachexia-relevant signaling. The upregulation of immune-associated transcripts in tumor cells, without host cell contamination, suggests that C26 cells acquire immune cell-like features, likely with the purpose to evade immune surveillance while simultaneously secreting cytokines that affect peripheral tissues.

Serum profiling from mice using cytokine array identified GDF15 and MMP9 as tumor-derived factors that were highly elevated during cachexia (Fig. 7). GDF15, a TGFβ superfamily cytokine, has been implicated in appetite suppression and systemic energy imbalance^54,55^ and has been implicated in the induction of SNAI1^56^. On the other hand, MMP9 is known as an ECM-remodeling protease, which plays an important role in ECM remodeling and controls cell adhesion and polarity^57,58^. MMP9 is also implicated in activation of latent TGFβ ligands^59^, and is shown to promote SNAI1 induction via SMAD3 phosphorylation^60^.

Although upregulation of Snai1 in muscle stem cells from cachectic mice and their impaired differentiation correlates with upregulation of tumor-derived GDF15 and MMP9, we cannot establish a direct relationship between Snai1 induction in muscle cells and GDF15/MMP9 from tumor cells with our current data. Interestingly, knockout of GDF15 in C26 tumors did not rescue muscle wasting, revealing redundancy from host or stromal sources such as macrophages and myofibers themselves (Fig. 8). This finding aligns with recent studies showing that GDF15 blockade mitigates anorexia but not muscle atrophy, highlighting its role as a systemic amplifier rather than a sole driver of cachexia.

Our findings converge on a model wherein C26 tumor cells transcriptionally reprogram themselves to evade immune surveillance. Progression of tumor growth leads to upregulation of tumor-derived GDF15 and MMP9 and concurrent upregulation of SNAI1 in the muscle stem cells. The net effect of tumor interaction with muscle stem cells is an altered metabolic state and stress response leading to impaired differentiation. By delineating how tumor-derived MMP9 and GDF15 converge on SNAI1 to repress myogenesis and destabilize the MuSC niche, we establish a mechanistic framework for tumor–muscle crosstalk and identify actionable targets for the prevention and reversal of muscle wasting in cancer.

## Materials

| Reagent or Resource | Source | Identifier |
| --- | --- | --- |
| Antibodies |  |  |
| PAX7 | DSHB | RRID: AB_528428 |
| MYOD1 (G1) | Santa Cruz | Cat# Sc-377460; RRID: AB_2813894 |
| Myogenin | DSHB | RRID: AB_2146602 |
| MF20 | DSHB | RRID: AB_2147781 |
| SNAI1 | Novus Biologicals | Cat# NBP2-27184; RRID: AB_2797904 |
| Phosphor-SMAD3 (S423/425) | Cell Signaling | Cat# 9520S; RRID: AB_2193207 |
| Anti-mouse ITGA7-Alexa647 | R&D Systems | Cat# FAB3518R; RRID:<br>AB_10973483 |
| Anti-mouse PE-CD11b | Invitrogen | Cat# 12-0112-82; RRID:<br>AB_2734869 |
| Anti-mouse PE-CD31 | BD Pharmigen | Cat# 553373; RRID: AB394819 |
| Anit-mouse PE-Sca1/Ly6A-E | BD Pharmigen | Cat# 553108; RRID: AB_394819 |
| Anti-mouse PE-CD45 | Invitrogen | Cat# 12-0451-82; RRID:<br>AB_465668 |
| Alexa Fluor 488 goat anti-rabbit<br>IgG | Invitrogen | A11008 |
| Alexa Fluor 594 goat anti-mouse<br>IgG2b | Invitrogen | A21145 |
| Alexa Fluor 594 goat anti-rabbit<br>IgG | Invitrogen | A11012 |
| Alexa Fluor 488 goat anti-mouse<br>IgG1 | Invitrogen | A21121 |
| Alexa Fluor 594 goat anti-mouse<br>IgG1 | Invitrogen | A21125 |
| Alexa Fluor 647 goat anti-rabbit<br>IgG | Invitrogen | A21245 |
| **Chemicals and Recombinant Proteins** |  |  |
| Fetal Bovine Serum | Wisent | 080-450 |
| 10% Goat Serum | Thermofisher | 50062Z |
| Horse Serum | Wisent | 065-150 |
| BSA | Sigma Aldrich | 9048-46-8 |
| Glycine | Bioshop | GLN001.5 |
| Sucrose | Millipore Sigma | S7903 |
| Triton-X100 | BioShop | TRX777.500 |
| Ham's F10 Nutrient Mix | Thermofisher | 11550043 |
| DMEM | Thermofisher | 11965118 |
| RPMI 1640 | Thermofisher | 11875093 |
| PBS 1X | Wisent | 311-013-CL |
| Penicillin-Streptomycin (10,000 U/mL) | Thermofisher | 15140122 |
| bFGF | Thermofisher | PHG0266 |
| Collagen from rat tail tendon | Millipore Sigma | 11179179001 |
| 2-Methylbutane | Millipore Sigma | 270342 |
| Frozen Section Compound | VWR | 95057-838 |
| Collagenase D | Millipore Sigma | 1108888201 |
| Dispase II | Millipore Sigma | 4942078001 |
| Hoechst | Thermofisher | 62249 |
| Paraformaldehyde | Electron Microscopy Sciences | 15714 |
| Prolong Gold Antifade Mountant with DNA Stain DAPI | Thermofisher | P36931 |
| Ampure XP beads | Beckman Coulter | A63882 |
| Cardiotoxin | Latoxan | L8102-1MG |
| L-Lysine-2HCl, 13C6, 15N2 | Thermofisher | 88432 |
| L-Arginine-<sup>13</sup>C<sub>6</sub>, <sup>15</sup>N<sub>4</sub> hydrochloride | Millipore Sigma | 608033 |
| Recombinant Mouse GDF-15 (CHO-expressed) Protein | R&D Systems | 10596-GD-025 |
| Recombinant Mouse MMP-9 Protein | R&D Systems | 909-MM-010 |
| **Commercial Kits** |  |  |
| Click-iT EdU Cell Proliferation Kit for Imaging Alexa Fluor 594 Dye | Thermofisher | C10339 |
| SMART-Seq HT Kit | Takara Bio | 634456 |
| Nextera XT DNA Library Preparation Kit | Illumina | FC-131-1024 |
| Nextera XT Index Kit | Illumina | FC-131-1001 |
| Tagment DNA Enzyme and Buffer (Small Kit) | Illumina | 20034210 |
| Proteome Profiler Mouse XL Cytokine Array | R&D System | ARY028 |

## Methods

### Mouse Strains and Animal Care

All mouse procedures conducted in this study were approved and performed in accordance with the guidelines set by the University of Ottawa’s Animal Care and Veterinary Services (ACVS) committee. Male and Female Balb/c mice were used in this study. All the mice used in the experiments are age and sex matched. All the mice that were used in experiments were 4 weeks old and 3-4 months old. Mice were maintained at 21°C with 20% humidity under a 14-hour light/10-hour dark cycle, food and water were provided *ad libitum*.

### Cell Culture

C26 adenocarcinoma cells were cultured with growth media (DMEM supplemented with 10% FBS, 1% penicillin/streptomycin) in an incubator at 37°C and 5% CO_2_. Cells were passaged at 70% confluency.

Primary myoblasts were cultured on collagen (Roche, 11179179001) coated plates with growth media (Ham’s F10 (Gibco, 11550043) supplemented with 20% FBS, 1% penicillin/streptomycin and 5 ng/mL of bFGF (Thermo Fisher, PHG2066)) in an incubator at 37°C and 5% CO_2_. Cells were passaged at 70% confluency.

### Primary Myoblast Differentiation

Cultured myoblasts were grown to 90% confluence and differentiated in DMEM supplemented with 5% horse serum (HS) and 1% penicillin/streptomycin for 5 days. After 3 days the media was replaced with fresh differentiation media for the duration of the assay.

### C26 Tumor Injection

Cultured C26 cells were trypsinized and the number of cells counted using Trypan Blue. The cells were suspended in 2% FBS/PBS. Mice were anesthetized with isoflurane and were injected twice, subcutaneously, with 1 million C26 cells each injection. Tumors were allowed to grow until the mice reached endpoint, or until the end of the experiment. Mice injected with Crispr C26 cells were given two injections of 2 million cells each.

### Generation of GFP C26 Adenocarcinoma Cells

Stable cell lines of C26 GFP expressing cells were generated as previously described^61^. Briefly, retrovirus particles containing the eGFP Plasmid were generated in Phoenix helper-free retrovirus producer lines using lipofectamine transfection. The viral supernatant was collected 48 h after transfection and subsequently applied to cultured C26 cells for 8 h. The cells were then washed twice with 1 × PBS and grown for 48 h in normal growth media. Cells were cultured in Puromycin Selection Media (DMEM supplemented with 10% FBS, 1% penicillin/streptomycin, and 2.5 μg/ml puromycin) for 1 week. The cells were then cultured in Low Puromycin Maintenance Media (DMEM supplemented with 10% FBS, 1% penicillin/streptomycin, and 1.25 μg/ml puromycin).

### Generation of GDF15 Crispr KO C26 Cells

CRISPR guide RNAs targeting exon 1 of Gdf15, this exon being common to all annotated isoforms, were designed^62^ and cloned into pLentiCRISPRv2_GFP^63^, such that guides did not share any potential off-target sequences. A control guide targeting Renilla luciferase was also cloned into the same vector. Guide sequences are given in Table 1. Lentiviral particles were generated from these vectors as previously described^64^ and were used to transduce C26 cells. Transduced cells expressing GFP were sorted using a Sony SH800 Cell Sorter. Seven days post-transduction, genomic DNA was isolated from sorted cells and was used as template in a PCR reaction with primer pair GCGATTGCGTCAGATTTCCG & CCGAATTAGCCTGGTCACCC. PCR products were purified and subjected to sequencing using Oxford Nanopore technology (Plasmidsaurus, OR) with data being analyzed for gene disruption efficiency using Crispresso2 software^65^. The populations generated with the most efficient guides (2 and 3) as well as the control population were again sorted using a Sony SH800 but this time to single cells to generate monoclonal populations. Genomic DNA from these populations was then isolated and used as template in PCR reactions with the same primer pair as before, with the PCR products being subjected to Sanger sequencing. Sanger reads were de-convolved using the Tide software suite^66^ to identify clones where out-of-frame knockout mutations had been introduced at the target site.

### Isolation of MuSCs by FACS

Muscle stem cells were isolated by Fluorescence Activated Cell Sorting (FACS) as previously described^53^. In brief, hindlimb muscles of the mice were dissected and mechanically minced. The minced muscles were transferred to a 15 mL conical tube and 5 mL of digestion buffer (F10 media, 10 U/mL Dispase II (Roche, 39307800), 2.4 U/mL Collagenase D (Roche, 11088882001) and 0.5mM CaCl_2_). Muscles were initially digested on a shaker for 20 minutes at 37 °C and 5% CO_2_. After the first round of digestion, the muscles were centrifuged for 10 seconds at 500g, and the supernatant was transferred to a fresh 50 mL conical tube containing 8 mL of FBS. The remaining muscle pellet was triturated and digested for a second time in an additional 5 mL of digestion buffer for 20 minutes. After the second digestion, the digested muscles were added to the same 50 mL conical tube containing the first digestion, and the entire mixture was filtered using a 40 µm cell strainer into a fresh 50 mL conical tube.

The filtered sample was centrifuged at 500g for 10 minutes at 4°C. The supernatant was removed and the pellet resuspended in 500 μL of FACS Buffer (2% FBS and 0.5mM EDTA in PBS) containing the antibody mixture (Alexa 647-ITGA7 (1:500) (R&D systems, FAB3518R, clone # 334908), PE-Cyanine7-VCAM-1 (1:500) (Invitrogen, 25-1061-82, clone # 429), PE-CD31(1:5000) (Invitrogen, 12-0311-82, clone # 390), PE-CD45 (1:5000) (Invitrogen, 12-0451-82, clone # 30F11), PE-CD11b (1:5000) (Invitrogen, 12-0112-82, clone # M1/70), PE-SCA1 (1:5000) (BD Biosciences, 553108, clone # D7) and Hoechst 33342 (Molecular Probes, H1399)). The samples were incubated for 20 minutes at RT with constant agitation and protected from light.

The samples were washed by adding 10 mL of FACS Buffer directly to the sample and centrifuging at 500g for 10 minutes. The supernatant was removed, and the cell pellet resuspended in 1 mL of FACS buffer. The samples were then filtered through a 40-µm cell strainer into a FACS compatible polypropylene round-bottom tube. The cells were sorted for the ITGA7^+^/VCAM1^+^/Hoechst^+^/CD31^−^/CD45^−^/CD11b^−^/SCA1^−^ population as described previously with a Sony SH800 Cell Sorter^53^.

### FACS Sorting C26 Cells from Whole Tumor

Mice that were injected with GFP C26 tumor cells were euthanized and the whole tumor dissected. The whole tumor was mechanically minced and digested in 5 mL of digestion buffer (F10 media, 10 U/mL Dispase II (Roche, 39307800), 2.4 U/mL Collagenase D (Roche, 11088882001) and 0.5mM CaCl_2_) for 15 minutes on a shaker at 37°C and 5% CO_2_. After, the digestion mix was centrifuged for 10 seconds at 500g and the supernatant transferred to a 50 mL conical tube containing 8 mL of FBS. The remaining tissue was triturated and another 5 mL of digestion buffer was added. The tissue was digested for 15 minutes on a shaker at 37°C and 5% CO_2_. The second digestion mixture was added to the 50 mL conical tube containing the first digestion and the sample was filtered using a 40 µm cell strainer into a fresh 50 mL conical tube. The sample was centrifuged at 500g for 10 minutes at 4°C. The cell pellet was resuspended in 1mL FACS Buffer (2% FBS and 0.5mM EDTA in PBS) containing DAPI and filtered through a 40 µm cell strainer into a 5 mL round bottom polypropylene tube. The GFP C26 cells were sorted for the GFP+/DAPI-population using a Sony SH800 Cell Sorter.

### Immunofluorescent Analysis of Cultured Cells

Cell culture media was removed, and cells were washed three times with PBS for 5 minutes at RT on a shaker. Cells were then fixed with 3.2% paraformaldehyde (PFA) (Electron Microscopy Sciences, 15714) in PBS for 5 minutes at RT and then washed three times in 1X PBS for 5 minutes. Cells were permeabilized for 15 minutes in 0.5% Triton-X and washed three times in 1X PBS for 5 minutes. Subsequently, cells were blocked in blocking buffer (0.1 M Glycine, 0.5% Bovine Serum Albumin (BSA) (Sigma, A8022), 0.3% Triton X-100 in PBS) for one hour at RT. Cells were incubated with primary antibodies diluted in the blocking buffer over night at 4 °C. The next day, the primary antibody solution was removed, and cells were washed three times with 0.1% Triton X-100 in PBS. After the washing step, secondary antibodies were diluted 1:400 in the blocking buffer and added to the cells for 1 hour at RT protected from light. Cells were washed three times with 0.1% Triton X-100 in PBS. Cells were mounted in mounting solution with DAPI and visualized under a microscope.

### Isolation and Culture of EDL Myofibers

Mice were sacrificed and the skin from the hindlimb was removed. The TA muscle was dissected out to expose the EDL muscle. The EDL was dissected by cutting the distal tendon and then the proximal tendon. The whole EDL was placed in an Eppendorf tube containing 800 μL of pre-heated (37 °C) myofiber digestion buffer (1000 U/ mL Collagenase (Sigma Aldrich, C0130) in unsupplemented DMEM). The EDL was placed in a tissue culture incubator with 5% CO_2_ and 37°C for 1 hour, with periodic mixing every 5-10 minutes.

A 6-well plate was coated with coating media containing 10% Horse Serum (HS) (Wisent, 065250) in DMEM for 30 minutes. The coating media was replaced with 2 mL of unsupplemented DMEM, and the plate was placed in the incubator to preheat the media. After the digestion was complete, a large bore glass pipette that is coated with coating media was used to transfer the EDL into the 6-well plate. Myofibers were mechanically dissociated by using the large bore glass pipetted and pipetting the EDL up and down. For the 0-hour time point, a subset of myofibers were transferred on a fresh 24-well plate and fixed immediately with 4% PFA. The remaining myofibers were incubated in the unsupplemented DMEM for 1 hour at 5% CO_2_ and 37°C. The DMEM was then removed and replaced with 2mL of pre-heated myofiber growth media (15% FBS, 1% chick embryo extract, 1% penicillin/streptomycin and 5 ng/mL of bFGF in DMEM).

### Immunofluorescence of EDL Myofibers

EDL myofibers were fixed at the selected times after isolation with 400 µL of 4% PFA in PBS for 5 minutes at RT and then transferred using a small bore glass pipette into a 24 well plate. Myofibers were washed with 400 µL of PBS three times. Myofibers were then permeabilized and blocked at the same time with 400 µL of blocking buffer (2% BSA, 6% HS, 10% GS, 1% Triton X-100, 0.1 M Glycine) for one hour at room temperature. Blocking buffer was then replaced with 200 µL of primary antibody master mix diluted in blocking buffer and incubated over night at 4 °C. After the primary antibody incubation, myofibers were washed three times with 400 µL of 0.1% Triton X-100 in PBS. Next, the samples were incubated with secondary antibody diluted in blocking buffer for 1 hour shielded from light at RT. Then, myofibers were washed three times with 400 µL of 0.1% Triton X-100 in PBS. Finally, myofibers were transferred onto a microscope slide, excess liquid was removed and mounting solution with DAPI was added.

### Immunofluorescent Staining of TA Muscle Cross Sections

TA muscle of the mice was dissected and fixed with 0.5% PFA in PBS for 2 hours at 4 °C. PFA was then removed and 20% sucrose in H_2_O was added to the muscle and incubated at 4 °C overnight. The TA muscle was then positioned in aluminum foil cups with OCT (Optimal Cutting Temperature) solution, and the muscle was frozen with liquid nitrogen-cooled isopentane. 9 μm thick cross-sections were cut using a cryostat and placed on microscope slides.

For immunofluorescence staining, cross-sections were first permeabilized with 0.5% Triton X for 15 minutes at RT followed by three 5 minute PBS washes. Cross-sections were blocked for 2 hours in blocking solution (3% BSA, 10% GS, 0.1M Glycine, in PBS) in a humid chamber at RT. Primary antibodies were diluted in the blocking solution and were then added and incubated overnight at 4 °C in the humid chamber. The next day, cross-sections were washed with 0.1% Triton X-100 in PBS for 5 minutes on a shaker, three times. After, secondary antibodies diluted 1:400 in the blocking solution were added and incubated for one hour at RT in the dark. Muscle sections were then washed three times with 0.1% Triton X-100 for 5 minutes on a shaker. Lastly, mounting solution with DAPI was added to the sections and cover slips were overlain on the slides to be visualized under the microscope.

### Cardiotoxin (CTX) Skeletal Muscle Injury

Skeletal muscle injury was caused by intramuscular injection of cardiotoxin (CTX) (Latoxan, L8102) into the hind limb of mice. 20 minutes prior to muscle injury, mice were subcutaneously injected with 100 µL of carprofen. Mice were anesthetized by isoflurane and intramuscularly injected with 50 µL of 10 µM cardiotoxin. Mice were sacrificed 7 days after the injury for further analysis to measure muscle regeneration.

### EchoMRI Readings

Mice were inserted into a size 40 tube and were pushed to the end of the tube using the plastic insert, leaving enough space for the length of the animal’s body. The tube containing the mouse was inserted into the EchoMRI and readings were taken with Primary Accumulation set to 1 and the Water Stage set to No. After readings, mice were returned to their home cage.

### Metabolic Cage

Metabolic readouts were measured using a Comprehensive Lab Animal Monitoring System (CLAMS) (Columbus Instruments). Mice were placed into individual live-in chambers within a larger environmental control unit. Temperature is maintained between 28 – 30°C as no bedding or nesting material is provided. A 12-hour light cycle is used for the duration of the experiment. Powdered food is provided *ad libitum* and is measured and recorded throughout the experiment. Animal movement, heat generation and respiration are all recorded throughout the experiment. Readings were taken every 20 minutes.

### Heavy Amino Acid Treatment

4-week-old male Balb/c mice were first treated with heavy Arginine (L-Arginine-^13^C_6_,^15^N_4_ hydrochloride (Millipore Sigma, 608033) for 1 week. The heavy Arginine was dissolved in the drinking water at 1 mg/mL and the mice were provided with 10 mL per mouse of water per day. The water was changed daily. After 7 days, the heavy Arginine water was replaced with normal drinking water for 2 days, then the mice were injected with C26 tumor cells as described above. The tumors were allowed to grow for 7 days with the mice being provided normal drinking water. After the first week of tumor growth, the normal water was replaced with water supplemented with 1 mg/mL of heavy Lysine (L-Lysine, 13C6, 15N2, Hydrochloride (Thermofisher, 88432). Each mouse was provided with 10 mL of water per day and the water was replaced daily. The mice were treated with the heavy Lysine water for 7 days and then euthanized. The whole muscle from the hindlimb and the tumor was isolated.

### Extraction and Digestion of Proteins for Total Proteomic Analyses

Proteins from powdered whole tissue were extracted in lysis buffer containing 5% sodium dodecyl sulfate (SDS), 100 mM TRIS pH 7.8. Samples were subsequently heated to 99°C for 10 minutes. The lysate was clarified by centrifugation 14,000 x g for 5 minutes. An aliquot corresponding to 10% of the total volume of lysate was diluted to <1% SDS and used for estimation of protein concentration by bicinchoninic acid assay (BCA). In the remaining sample, protein disulfide bonds were reduced by the addition of tris(2-carboxyethyl) phosphine (TCEP) to a final concentration of 20 mM and incubated at 60°C for 30 minutes. Free cysteines were alkylated using iodoacetamide at a final concentration of 30 mM and subsequent incubation at 37°C for 30 minutes in the dark. An equivalent of 10 μg of total protein was used for proteolytic digestion using suspension trapping (STRAP). In brief, proteins were acidified through the addition of phosphoric acid to a final concentration of 1.3% v/v. The sample was subsequently diluted 6-fold in STRAP loading buffer (9:1 methanol: water in 100 mM TRIS, pH 7.8) and loaded onto an S-TRAP Micro cartridge (Protifi LLC, Huntington NY) and spun at 4000 x g for 2 minutes. Samples were washed three times using 150 μL of STRAP loading buffer. Proteins were then proteolytically digested using trypsin at a 1:10 enzyme to substrate ratio for 2 hours at 47°C. Peptides were sequentially eluted in 50 mM ammonium bicarbonate, 0.1% formic acid in water, and 50% acetonitrile. Peptides were vacuum concentrated and reconstituted in 0.1% formic acid (FA) prior to analysis by LC-MS/MS.

### LC-MS/MS Acquisition and Data Analysis

Peptide containing samples were analyzed by data dependent acquisition (DDA-PASEF) using anEvosep One LC system and a Bruker timsTOF HT mass spectrometer. Samples (200ng) were loaded onto Evotips according to the vendor protocol (Evosep). Peptides were separated using the 30 samples per day method and an EV1137 analytical column (Evosep) set to 45°C. Full MS scans were acquired from m/z 100-1,700, between an ion mobility window of 0.6 – 1.6 1/k0 thus filtering for primarily 2+ and 3+ precursor charge states. The 10 most intense ions rPASEF ramp and accumulation times were set to 100 ms and fragmented using a normalized collision energy scaled based on ion mobility – from 20 eV at 0.6 1/k0 to 59 eV at 1.6 1/k0. and the dynamic exclusion was set to 40 s. DDA-PASEF MS raw data was processed with Fragpipe version 22 (https://github.com/Nesvilab/FragPipe)^67^ with database searches against a FASTA file containing all reviewed canonical sequences downloaded from UniProt. The enzyme specificity was set to trypsin with a maximum of 2 missed cleavages. Carbamidomethylation of cysteine was set as fixed modification and oxidation of methionine as variable modification. The precursor and product ion mass tolerances were set to 20 ppm. Proteins were quantified by label free quantitation using IonQuant with default settings. In order to estimate the relative amount of a particular protein within the sample we calculated and ranked proteins based on their normalized spectral abundance factor (NSAF) values (https://doi.org/10.1021/pr060161n). Hierarchical clustering was conducted using the normalized protein LFQ abundances as inputs for Instant Clue (http://www.instantclue.uni-koeln.de/) and FragPipe-analyst^68^.

### Preparation of SMART RNA-Seq library MuSCs and C26 cells

SMART-Seq was performed as previously described with the use of the SMART-Seq HT Kit (TAKARA, 634456)^53^. A 10X SMART-Seq lysis buffer working solution was prepared (95% 10X SMART-Seq lysis buffer, 5% RNase Inhibitor). 1000 MuSCs or C26 tumor cells were sorted by FACS into 10 µL of 1X-SMART Lysis Buffer (1 µL of 10X SMART-Seq lysis buffer working solution + 9 µL ddH_2_O). After, 1 µL of 3’ SMART-Seq CDS primer II A was added, and then incubated for 3 minutes at 72 °C in a thermocycler. The template switching master mix (0.7 µL of nuclease free water, 8 µL of one-step buffer, 1 µL SMART-Seq HT oligonucleotide, 0.5 µL RNase inhibitors, 0.3 µL of SeqAMP DNA polymerase and 2 µL of SMARTscribe reverse transcriptase) was prepared and 12.5 µL was added. The cDNA was amplified for 11 cycles as previously described^53^. cDNA was purified with AmpureXP at a 1:1 ratio (v/v). The beads were washed twice with 200 µL of 80% ethanol and the DNA was then eluted with 18 µL of elution buffer. The cDNA was then quantified with a Quant-IT Picogreen dsDNA Assay Kit (Invitrogen, P7589).

Sequencing ready libraries were prepared with Nextera XT Library Prep Kit (Illumina, FC-131-1024) as previously described^53^. In brief, 250 pg of cDNA in a volume of 1.25 µL was transferred to a fresh microtube. 2.5 µL of Tagment DNA (TD) buffer and 1.25 µL Amplicon Tagment Mix (ATM) Buffer were added for a total of 5 µL volume. The cDNA was incubated at 55 °C for 5 minutes in a thermocycler. Immediately after, 1.25 µL of NT Buffer was added and incubated at least 5 minutes at RT. Next, 3.75 µL of NPM PCR master mix, 1.25 µL each of an i7 and an i5 index from the Nextera XT index adaptor kit (Illumina, FC-131–2001) were added for a total of 12.5 µL. The samples were then PCR amplified for 12 cycles as previously described^53^. Lastly, the PCR amplified libraries were purified, and size selected with 0.85X ratio (v/v) AmpureXP beads and were sequenced.

### ATAC-Seq of MuSCs

OMNI ATAC-Seq was performed on 5000 freshly sorted MuSCs from 4-month-old healthy control and tumor bearing female mice, as previously described^36^. MuSCs were FACS sorted into 30 µL of ATAC lysis buffer (10 mM Tris-HCl, 10 mM NaCl, 3 mM MgCl_2_, 0.1% Tween-20, 0.1% NP-40 and 0.01% Digitonin (Promega, G9441)) and incubated on ice for 5 minutes and then at RT for 3 minutes. Cells were then washed with 100 µL of 10 mM Tris-HCl (pH 7.5), 10 Mm NaCl, 3 mM MgCl_2_ and 0.1% Tween-20, and centrifuged at 800g for 10 minutes. The supernatant was removed and the pellet was resuspended in 10 µL of transposition mixture (5 µL TD buffer, 3.2 µL PBS, 0.89 µL Tn5 (Illumina, 20034197), 0.1% Tween-20, 0.01% Digitonin and 0.75 µL nuclease free water) and incubated at 37 °C for 25 minutes. DNA Purification was performed using a QIAquick PCR Purification Kit (Qiagen, 28,104) according to the manufacturer’s guidelines and eluted in 20 µL of elution buffer.

Indices were incorporated by adding 30 μL of the PCR reaction mix (10 μL Q5 buffer, 10 μL Q5 enhancer, 1 μL dNTPs, 2.5 μL i7 index, 2.5 μL i5 index, 3.5 μL nuclease free water, 0.5 μL Q5 High Fidelity DNA polymerase). PCR amplification was performed using the previously described conditions for 12 cycles^36^. The libraries were size selected and purified with Ampure XP beads at a 1:0.85 (v:v) ratio. Samples were sequenced with NovaSeq6000 Sprime, Paired End (PE) 150bp.

### Single myofiber RNA-Seq

Single myofiber RNA-Seq (smfRNA-Seq) was performed on EDL myofibers isolated from 4-month-old male and female, healthy control and tumor bearing mice. The procedure was performed as previously described using the SMART-Seq HT Kit and Nextera XT Library Prep Kit^18^. After the dissociation of the myofibers, a coated small-bore glass pipette was used to transfer the myofibers into a 6-well plate containing PBS and a single myofiber was selected and transferred into a 0.2 mL PCR tube. PCR tube was then visualized under the microscope to ensure the selection of one myofiber. The excess PBS was then removed while visualizing under the microscope. 1X SMART lysis buffer was prepared as described in the section, SMART RNA-Seq library preparation of MuSCs. Then 10 µL of the 1X Lysis Buffer was added to the PCR tube containing the myofiber. The fiber was pipetted up and down and lysed on ice for 5 minutes, with vortexing as needed. The sample was then spun down to remove the residual fiber pieces. While visualizing under the microscope, the supernatant was transferred to a new 0.2 mL PCR tube. The cDNA library preparation was performed as described in the above section, SMART RNA-Seq library preparation of MuSCs except for SMART cDNA amplification being 12 cycles. The rest of the library preparation using the Nextera XT library prep kit was performed as described in the previous sections. n=3 RNA-Seq of myofibers was performed to have enough samples for downstream analysis.

### EdU Incorporation Assay

EdU incorporation assay was performed using the Click-It Plus EdU Alexa Fluor 555 Imaging Kit (Invitrogen, C10638) according to the manufacturer’s guidelines. Cells were seeded into chamber slides prior to EdU Assay. Briefly, 10 µM of EdU was added to the growth media of cultured cells for 12 hours. Following the incubation period, media was removed, and the cells were washed with PBS three times. The cells were then fixed with 3.2% PFA and EdU staining was performed as described by the manufacturer.

### RNA-seq Analysis

Single-end RNA-seq reads were quantified with Kallisto v0.46.1 (parameters: –single –fragment-length 200 –sd 20)^69^. Ensembl gene annotations, release 96, were used to create an index for Kallisto^70^. Transcript-level abundance estimates were collapsed to the gene level using ‘‘tximport’’^69^. Pseudoalignments constructed by kallisto were converted to bigwigs using bamCoverage (version 3.5.1)^71^. Counts were normalized to RPM.

Differentially expressed genes were identify using DESeq2^72^. P values were adjusted for multiple testing using the Independent Hypothesis Weighting procedure^73^.

Gene set enrichment analysis was performed using fgsea, the log fold changes, and the Hallmark gene sets collection from Molecular Signatures Database (MsigDB) (https://doi.org/10.1101/060012)^74,75^. Library size normalized gene counts were visualized with ComplexHeatmap^76^.

### ATAC-Seq Analysis

Adapters were removed from raw ATAC-seq reads using Trimmomatic (version 0.39, parameters: LEADING:10 TRAILING:10 MINLEN:50)^77^. Paired reads were aligned to mm10 using BWA-MEM (version 0.7.17). Samtools was used to index and filter (version 1.16.1, mapping quality > 30)^78^. Duplicated reads were removed using Picard (version 3.1.0). ATAC-seq peaks were called with MACS3 (version 3.0.1, parameters: callpeak, –g mm, –q 0.05)^79^.

The summits from each sample are pooled and summits closer to 250 bps are merged. Then, the coordinates are extended 250 bps from the center. Read counts are retrieved from the bam files using featureCounts (version: 2.1.1, –p –F SAF ––fracOverlap 0.2)^80^.

Differentially bound peaks were identify using DESeq2^72^. P values were adjusted for multiple testing using the Independent Hypothesis Weighting procedure^73^. Peaks were associated to the genes with the closest TSS. Gene set enrichment analysis was performed as described before.

ATAC-seq samples were merged by condition (control and cachexia). Correction of Tn5 insertion bias and calculation of footprint scores were performed using TOBIAS (subcommands: ATACorrect and FootprintScores, with defaults options)^81^. Differentially bound motifs were estimated using TOBIAS (subcommand: BINDetect, defaults options) the Jaspar database (collection: core, vertebrates, non redundant)^82^.

### Quantification and statistical analysis

Statistical analyses were performed using Prism10 (Graphpad). All data is presented as mean ± SD. When two independent groups were compared an unpaired two-tailed t-tests was used. When multiple groups were compared, analysis was done using a 2-way ANOVA. Exact p values are found within the relevant figure panel and a p-value of < 0.05 was considered to be significant. Sample sizes for each experiment are described in the corresponding figure legend.

## Data Availability

All sequencing data generated in this study were deposited to the Gene Expression Omnibus under the super series GSE310038. Individual accession numbers for the RNA-Seq and ATAC-Seq data are GSE309918 and GSE309918 respectively.

## Acknowledgements

We thank Dr. Vera Tang and the Flow Core facility at the University of Ottawa for their help with Fluorescence Activated Cell Sorting and Flow Cytometry analysis. We would also like to thank the Animal Behaviour and Physiology Core, funded by the University of Ottawa Faculty of Medicine, for their assistance with the EchoMRI and metabolic cage data. We thank Mrs. Jothi Krishnamoorthy for excellent technical assistance in some experiments. Cell lines with gene disruptions were generated at the Genomic Engineering and Molecular Biology (GEMb) core facility in the Faculty of Medicine at the University of Ottawa (RRID: SCR_022954). The GEMb facility is supported by CFI-36490, CFI-37607 and CFI-36940. LentiCRISPRv2GFP was a gift from David Feldser (Addgene plasmid # 82416; http://n2t.net/addgene:82416; RRID:Addgene_82416). This work was supported by a Molecular and Regenerative Medicine Pilot Grant Program from the Lady Davis Institute, Jewish General Hospital to V.D.S and A.E.K, as well as from Canadian Institutes of Health Research [CIHR; PJT-178173 and PJT-168864]to A.E.K and Canadian Institutes of Health Research [PJT-190082] to V.D.S.

## Author Contributions

Conceptualization: D.M.B, A.E.K and V.D.S.; Methodology: D.M.B, K.S, V.R, F.L, A.H.C, D.Q, S.H and H.H.K; Visualization: D.M.B, V.D.S, and A.H.C.; Investigation: D.M.B, A.H.C, and V.D.S; Computational Analysis: A.H.C; Writing manuscript: D.M.B and V.D.S; Supervision: V.D.S, H.N, and C.B.; Funding Acquisition: V.D.S.; Resources: V.D.S, G.B, M.A.R, Y.Y, A.J.A; Reviewing/editing: D.M.B, K.S, F.L and V.D.S.

