## Supplemental Information for "A Multiomic Analysis of Cachectic Mice Reveals Cancer Driven Suppression of Muscle Stem Cell Differentiation"

**Extended Data Figures 1-8**

Extended Data 1

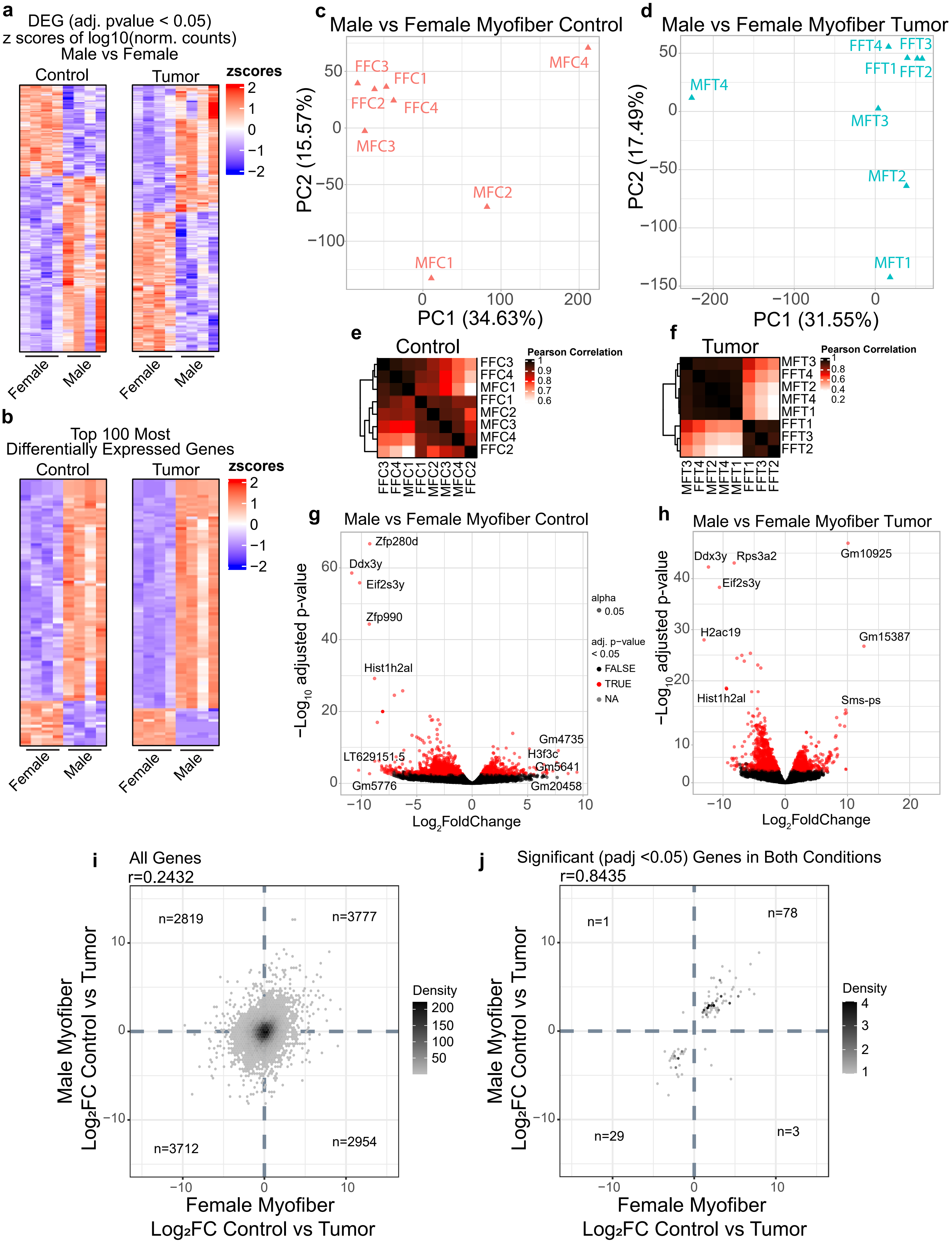

**Extended Data Figure 1: Transcriptional variation between male and female myofibers**

**a.** Heatmap of the differentially expressed genes (adj. p-value $\leq$ 0.05) between male and female myofibers from tumor bearing (left) and healthy control (right) mice. **b.** Heatmap of the top 100 most differentially expressed genes (adj. p-value $\leq$ 0.05) between male and female myofibers from tumor bearing (left) and healthy control (right) mice. **c.** PCA plot of sequenced single myofibers from healthy 3-month-old male and female mice. **d.** PCA plot of sequenced single myofibers from tumor bearing 3-month-old male and female mice. **e.** Pearson correlation between male and female myofibers isolated from healthy controls. **f.** Pearson correlation between male and female myofibers isolated from tumor bearing mice.**g.** Volcano plot of male vs female myofibers isolated from healthy controls. **h.** Volcano plot of male vs female myofibers isolated from tumor bearing mice. **i.** Scatterplot of the effect of cachexia on transcript levels between male and female myofibers. Comparison between the LFC of all genes of healthy controls vs cachexia in male with the LFC of female healthy controls vs cachexia. **j.** Scatterplot comparing the LFC of healthy controls vs cachexia in male with the LFC of female healthy controls vs cachexia. Only genes that are significantly different (adjusted p-value $\leq$ 0.05) in both groups were plotted.

Extended Data 2

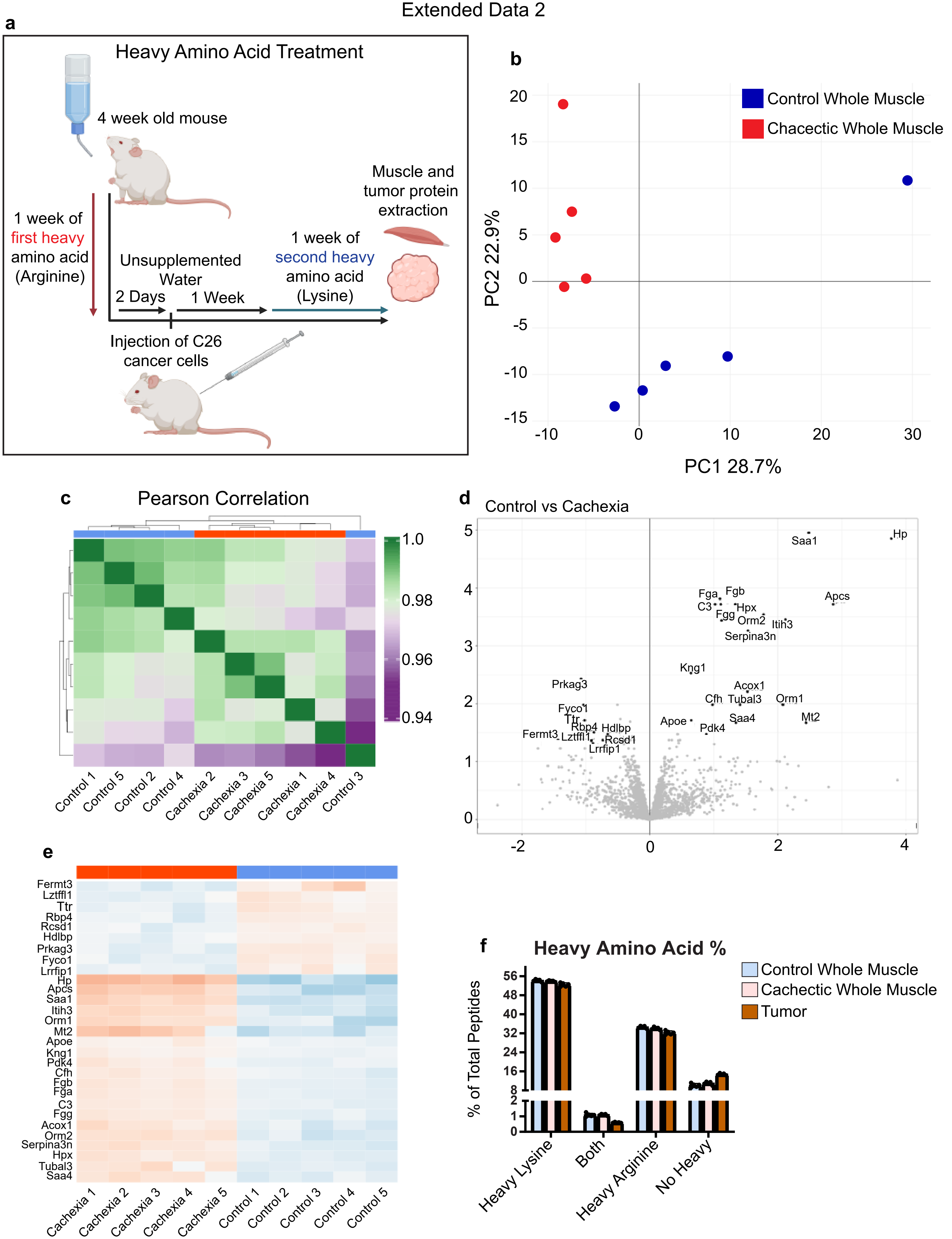

**Extended Data Figure 2: Muscle proteome is significantly altered during cachexia**

**a.** PCA plot of control and cachectic whole muscle. **b.** Pearson correlation between control and cachectic whole muscle samples. **c.** Heatmap of all the significantly different (LFC $\geq$ 1.5 and adjusted p-value $\leq$ 0.05) proteins between control and cachectic muscle. **d.** Volcano plot of the LFC in protein expression between control and cachectic muscle. Significantly different proteins are labelled. **e.** Bar plot of the percentage of peptides containing heavy lysine, heavy arginine, both, or none in the control and cachectic whole muscle and tumor whole tissue samples.

Extended Data 3

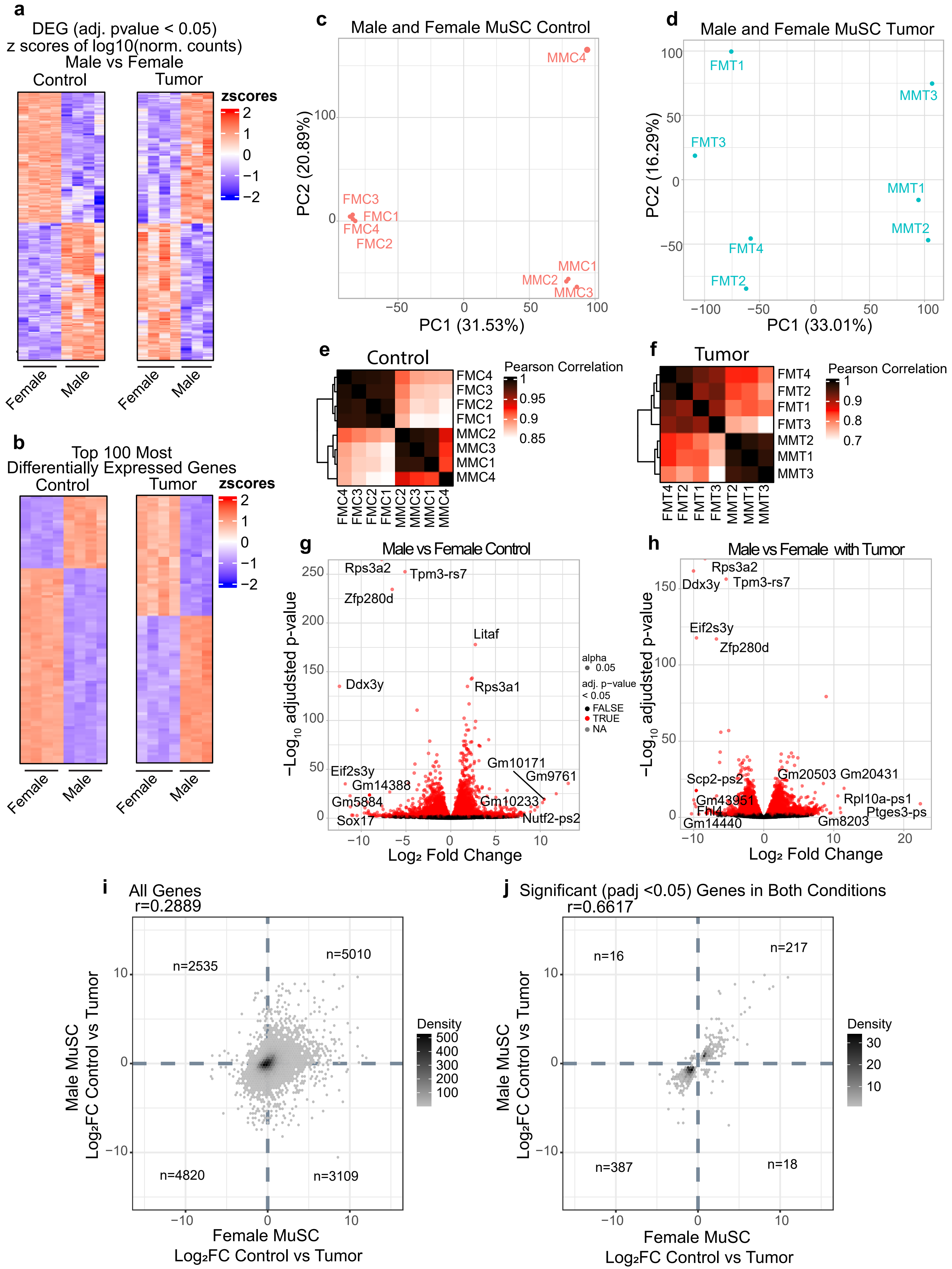

**Extended Data Figure 3: Male and female MuSCs have a distinct transcriptomic profile but are similarly affected by cachexia.**

**a.** Heatmap of the differentially expressed genes (adj. p-value $\leq$ 0.05) between male and female MuSCs from tumor bearing (left) and healthy control (right) mice. **b.** Heatmap of the top 100 most differentially expressed genes (adj. p-value $\leq$ 0.05) between male and female MuSCs from tumor bearing (left) and healthy control (right) mice. **c.** PCA plot of sequenced MuSCs from healthy 3-month-old male and female mice. **d.** PCA plot of sequenced MuSCs from tumor bearing 3-month-old male and female mice. **e.** Pearson correlation between male and female MuSCs isolated from healthy controls. **f.** Pearson correlation between male and female MuSCs isolated from tumor bearing mice. **g.** Volcano plot of male vs female MuSCs isolated from healthy controls. **h.** Volcano plot of male vs female MuSCs isolated from tumor bearing mice. **i.** Scatterplot of the effect of cachexia on transcript levels between male and female MuSCs. Comparison between the LFC of all genes of healthy controls vs cachexia in male with the LFC of female healthy controls vs cachexia. **j.** Scatterplot comparing the LFC of healthy controls vs cachexia in male with the LFC of female healthy controls vs cachexia. Only genes that are significantly different (adjusted p-value $\leq$ 0.05) in both groups were plotted.

Extended Data 4

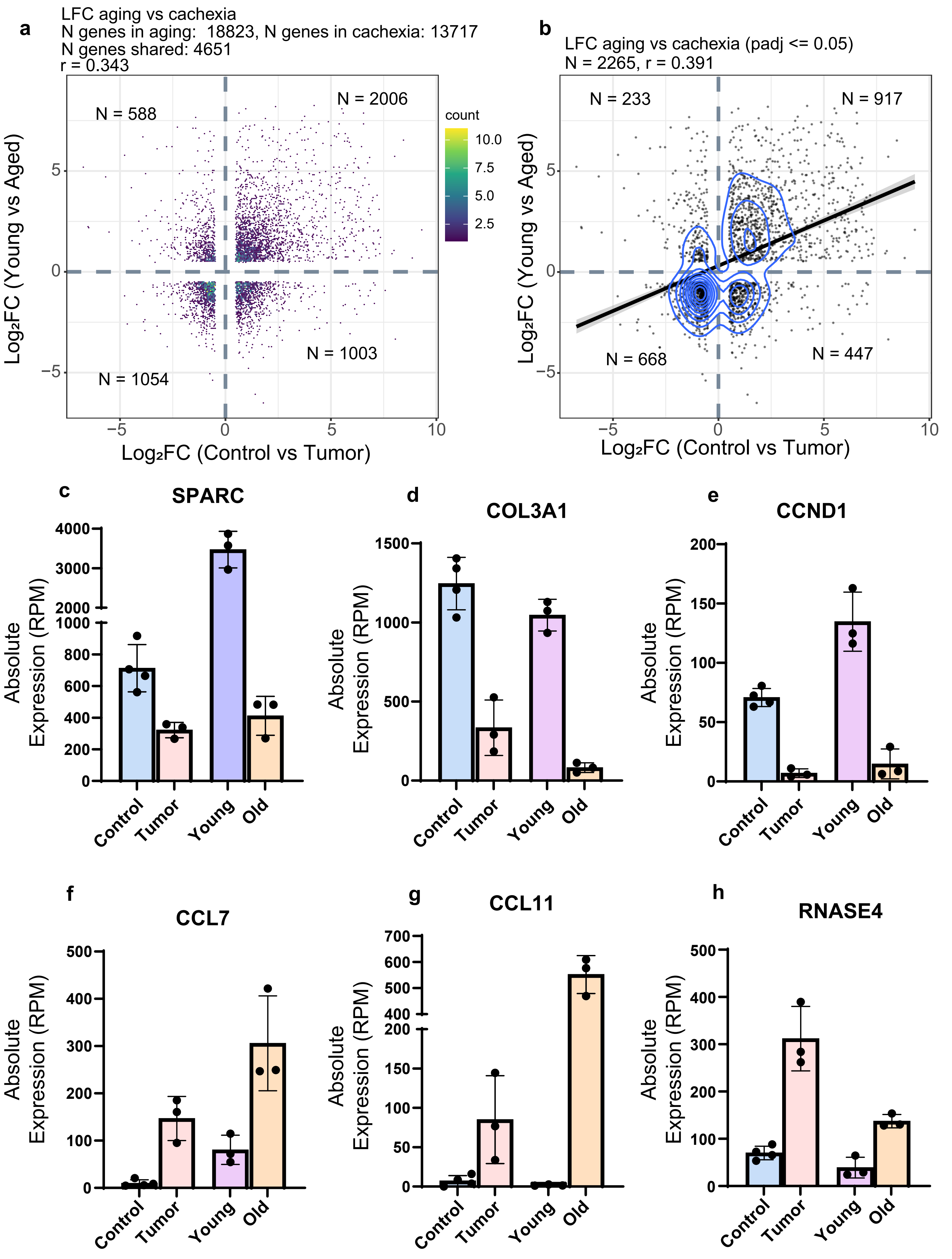

**Extended Data Figure 4: Cachexia and aging have similar effects on MuSC transcriptome**

**a.** Scatter plot comparing LFC of healthy control male MuSCs vs cachectic male MuSCs and the LFC of young male MuSCs and aged male MuSCs. The young and aged MuSC RNA-Seq data is publicly available and was retrieved from GSE171997. Only genes that had an LFC $\geq$ 0.5 in both conditions were plotted. **b.** Scatter plot comparing LFC of healthy control male MuSCs vs cachectic male MuSCs and the LFC of young male MuSCs and aged male MuSCs. Only significant genes were plotted (LFC $\geq$ 0.5 and adjusted p-value $\leq$ 0.05). Bar graphs of the absolute expression (RPM) of **c.** Sparc, **d.** Col3a1, **e.** Ccnd1, **f.** Ccl7, **g.** Ccl11, **h.** RNASE4 between male healthy controls, tumor bearing, young and aged MuSCs.

Extended Data 5

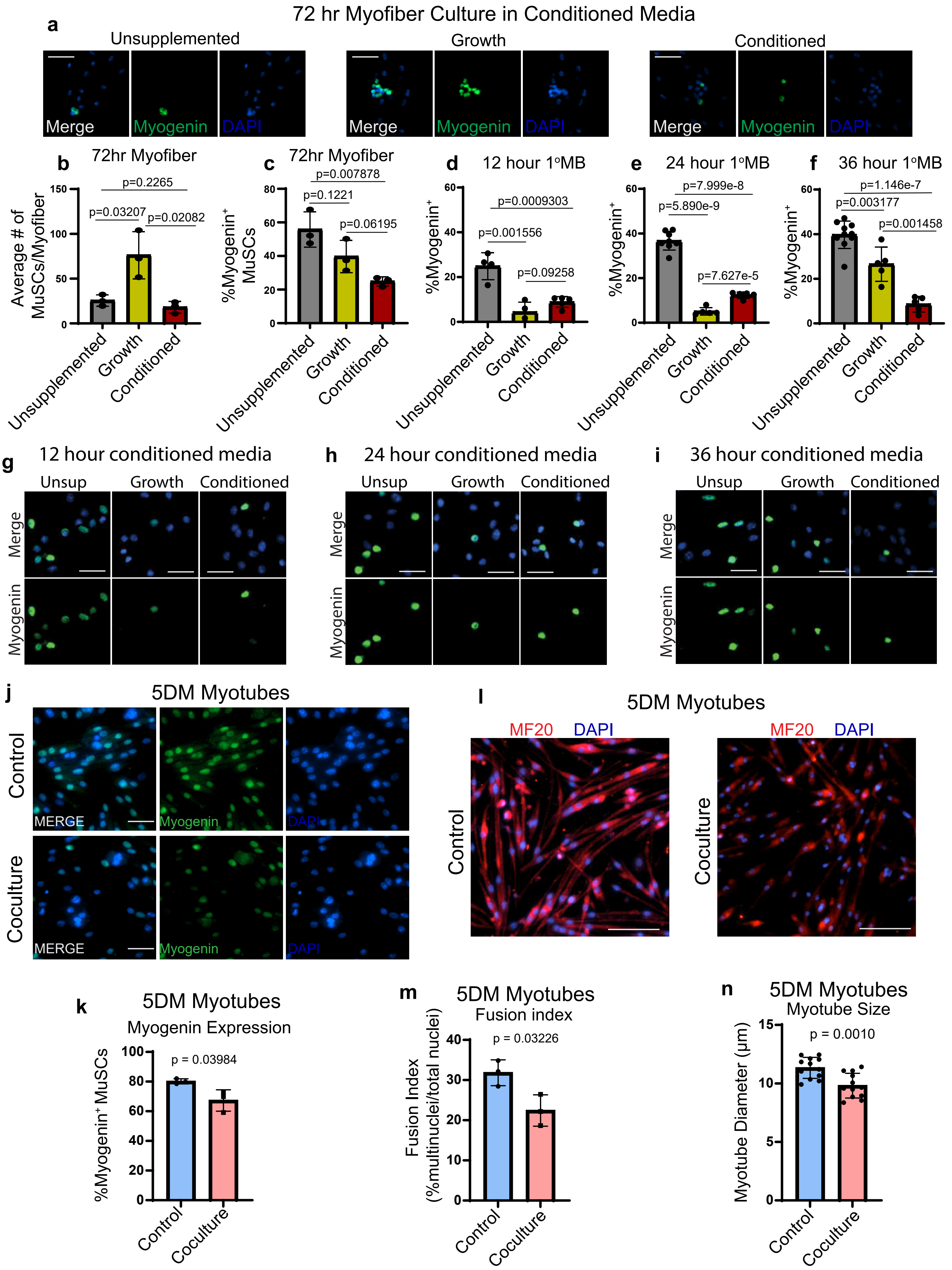

**Extended Data Figure 5: C26 cells reduce muscle stem cell growth and differentiation**

**a.** Representative pictures of EDL myofibers cultured for 72 hours in Unsupplemented Growth Media (15% FBS, 1% P/S, - CEE, - bFGF), Normal Growth Media (15% FBS, 1% P/S, 1% CEE, 5 ng/µL of bFGF) and C26 Conditioned Media (Normal Growth Media cultured for 48 hours with C26 cells). Myofibers were stained for Myogenin. Scale bar = 40 µm. **b.** Quantification of the total number of MuSCs per myofiber, cultured for 72 hours. n = 3 biological replicates. **c.** Quantification of the percentage of Myogenin positive MuSCs per myofiber, cultured for 72 hours. n = 3 biological replicates. Percentage of Myogenin positive primary myoblasts cultured in Unsupplemented, Normal Growth, and C26 Conditioned media for **d.** 12 hours, **e.** 24 hours, and **f.** 36 hours. n = 3 biological replicates. Representative images of primary myoblasts cultured in Unsupplemented Growth Media (20% FBS, 1% P/S), Normal Growth (20% FBS, 1% P/S, 5 ng/µL of bFGF), and C26 Conditioned Media (Normal Growth Media cultured for 48 hours with C26 cells) for **g.** 12 hours, **h.** 24 hours, **i.** and 36 hours. Primary myoblasts were stained for Myogenin. Scale bar = 30 µm. **J.** Representative picture of primary myotubes differentiated for five days, with or without coculture with C26 cells. Cells were stained for Myogenin. Scale bar = 40 µm. **k.** Quantification of the percentage of Myogenin positive nuclei in 5DM myotubes, with or without C26 coculture. n = 3 biological replicates. **l.** Representative pictures of primary myotubes differentiated for 5 days, with or without C26 coculture. Cells were stained for Myosin Heavy Chain. Scale bar = 60 µm. **m.** Quantification of the fusion index of 5DM primary myotubes, with or without C26 coculture. n = 3 biological replicates. **n.** Quantification of the average myotube diameter, defined as the widest region across the myotube, of 5DM primary myotubes with or without C2 coculture. n = 3 biological replicates, 4 fields of view per replicate.

Extended Data 6

**a** Number of Differentially Expressed Genes

| Threshold | Adjusted p-value ≤0.1 | | | Adjusted p-value ≤0.01 | | | Adjusted p-value ≤0.001 | | |
|---|---|---|---|---|---|---|---|---|---|
| Comparison | In vitro vs Female | In vitro vs Male | Female vs Male | In vitro vs Female | In vitro vs Male | Female vs Male | In vitro vs Female | In vitro vs Male | Female vs Male |
| Number of Upregulated DEGs | 7880 | 6344 | 200 | 6017 | 4652 | 38 | 4708 | 3410 | 11 |
| Number of Downregulated DEGs | 3155 | 4137 | 272 | 1567 | 2571 | 48 | 907 | 1779 | 20 |

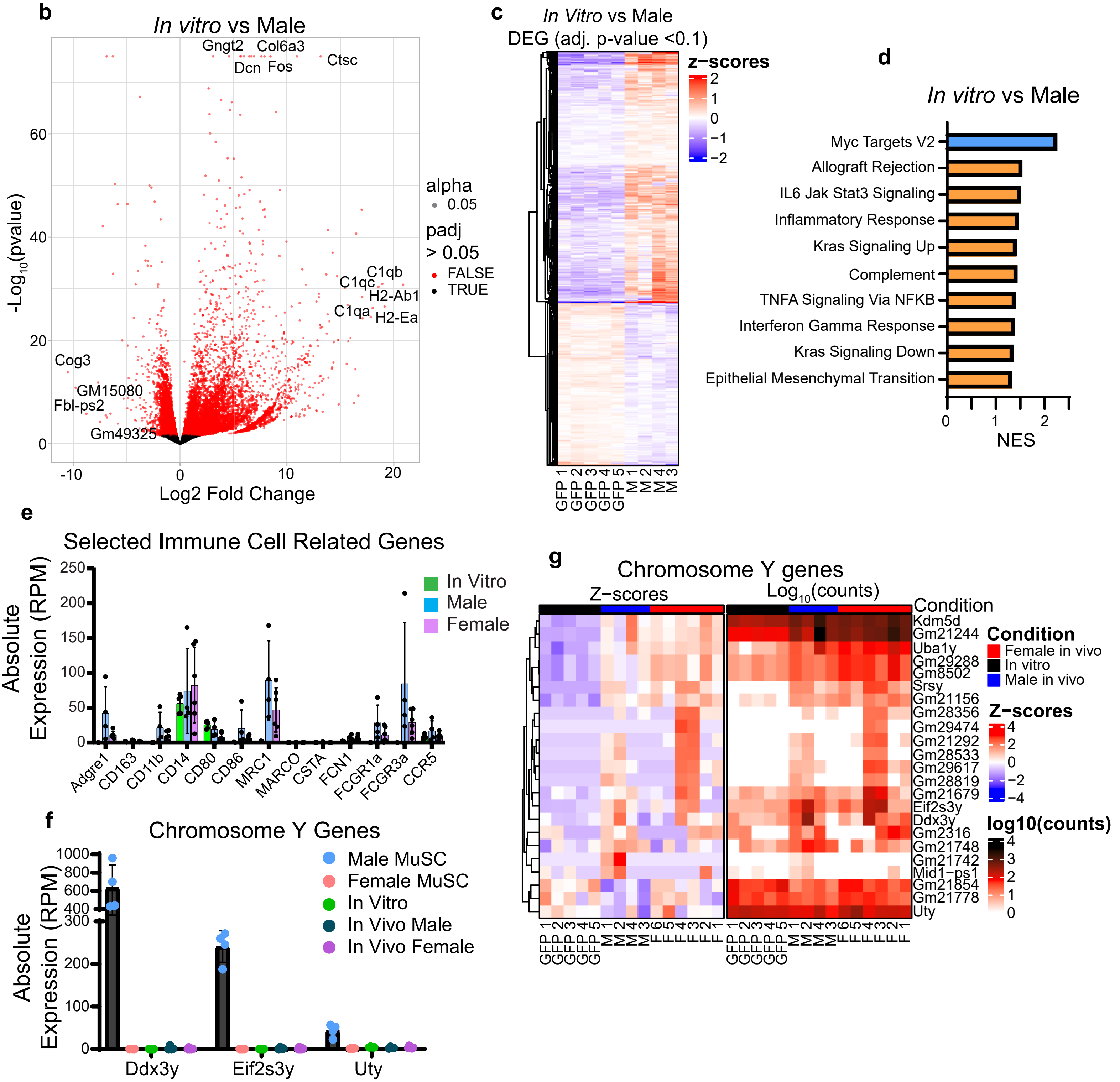

**Extended Data 6: Sex of the host does not affect transcriptional adaptation of C26 cancer cells**

**a.** Table of the number of differentially expressed genes between *In vitro* C26 vs *In vivo* in Female, *In vitro* C26 vs *In vivo* in Male, and *In vivo* in Female vs *In vivo* in Male. **b.** Volcano plot comparing gene expression between *In vitro* C26 and *In vivo* in Male. **c.** Heatmap of the differentially expressed (adjusted p-value $\leq$ 0.1) between *In vitro* C26 vs *In vivo* in Male. **d.** Top ten most significantly enriched Hallmark pathways between *In vitro* C26 vs *In vivo* in Male. Blue bars indicate a pathway is downregulated, orange bars indicate a pathway is upregulated. **e.** Bar plot indicating the absolute expression (RPM) of selected immune cell related genes in *In vitro* C26, *In vivo* in Female, and *In vivo* in Male RNA-Seq samples. **f.** Bar plot of the absolute expression (RPM) of selected Y chromosome genes in Male MuSCs, Female MuSCs, *In vitro* C26, *In vivo* in Female, and *In vivo* in Male RNA-Seq samples. **g.** Heatmap of the Y chromosome gene expression in *In vitro* C26, *In vivo* in Female, and *In vivo* in Male RNA-Seq samples.

Extended Data 7

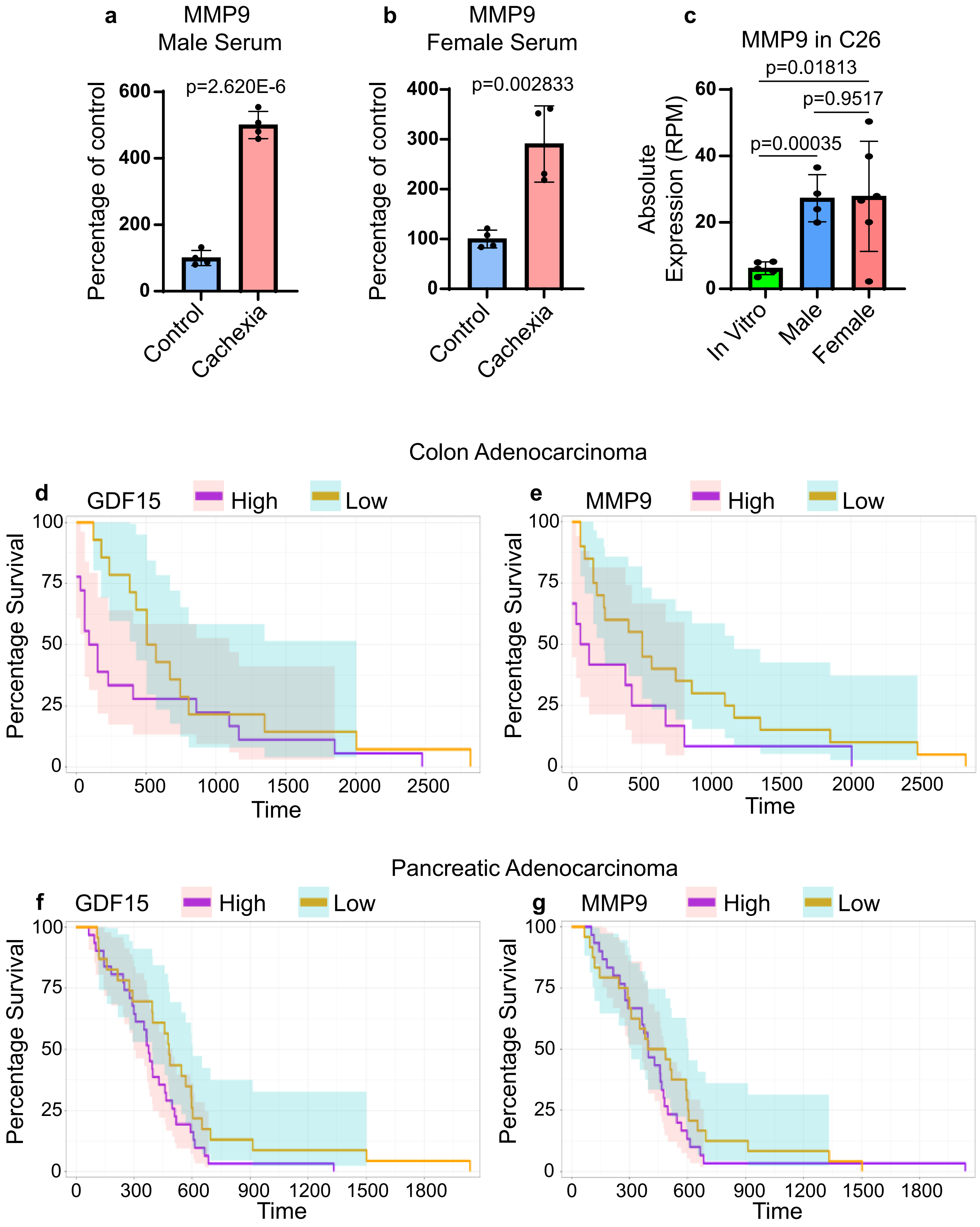

**Extended Data 7: Cancer patients with high expression of GDF15 and MMP9 have a worse prognosis**

**a.** Quantification of MMP9 levels in the serum of male healthy control and tumor bearing mice. **b.** Quantification of MMP9 levels in the serum of male healthy control and tumor bearing mice. **c.** Expression of MMP9 in C26 cells. **d.** Kaplan Meier curve of colon adenocarcinoma patients with high and low expression of GDF15 in the tumor. GDF15 high n = 22; GDF15 low n = 14. **e.** Kaplan Meier curve of colon adenocarcinoma patients with high and low expression of MMP9 in the tumor. MMP9 high n = 16; MMP9 low n = 20. **f.** Kaplan Meier survival curve of pancreatic adenocarcinoma patients with high and low expression of GDF15 in the tumor. GDF15 high n = 31; GDF15 low n = 23. Data was retrieved from TCGA and only patients that did not undergo chemotherapy or surgery were selected. **g.** Kaplan Meier survival curve of pancreatic adenocarcinoma patients with high and low expression of MMP9 in the tumor. MMP9 high n = 30; MMP9 low n = 24. Data was retrieved from TCGA and only patients that did not undergo chemotherapy or surgery were selected.

**Extended Data 8: MMP9 treatment increases SNAI1 expression in primary myoblasts**

**a.** Representative images of primary myoblasts treated with MMP9 and stained for MyoD1, SNAI1, and Myogenin. Scale bar = 30 um. n = 3 biological replicates. **b.** Percentage of the high SNAI1 expressing primary myoblasts treated with MMP9. **c.** Percentage of Myogenin positive primary myoblast treated with MMP9. **d.** Representative image of isolated EDL myofibers cultured for 36 hours before being treated with MMP9 for 1 hour. n = 3 biological replicates. Myofibers were stained with MyoD and phosphor-SMAD3. **e.** Percentage of high phosphor-SMAD3 expressing MuSCs per myofiber. **f.** Representative image of isolated EDL myofibers cultured for 60 hours and stained for MyoD and Myogenin. n = 3 biological replicates. **g.** quantification of the number of MuSCs per myofiber after 60 hours in culture. **h.** Quantification of the percentage of Myogenin positive MuSCs per myofiber.

Extended Data 8

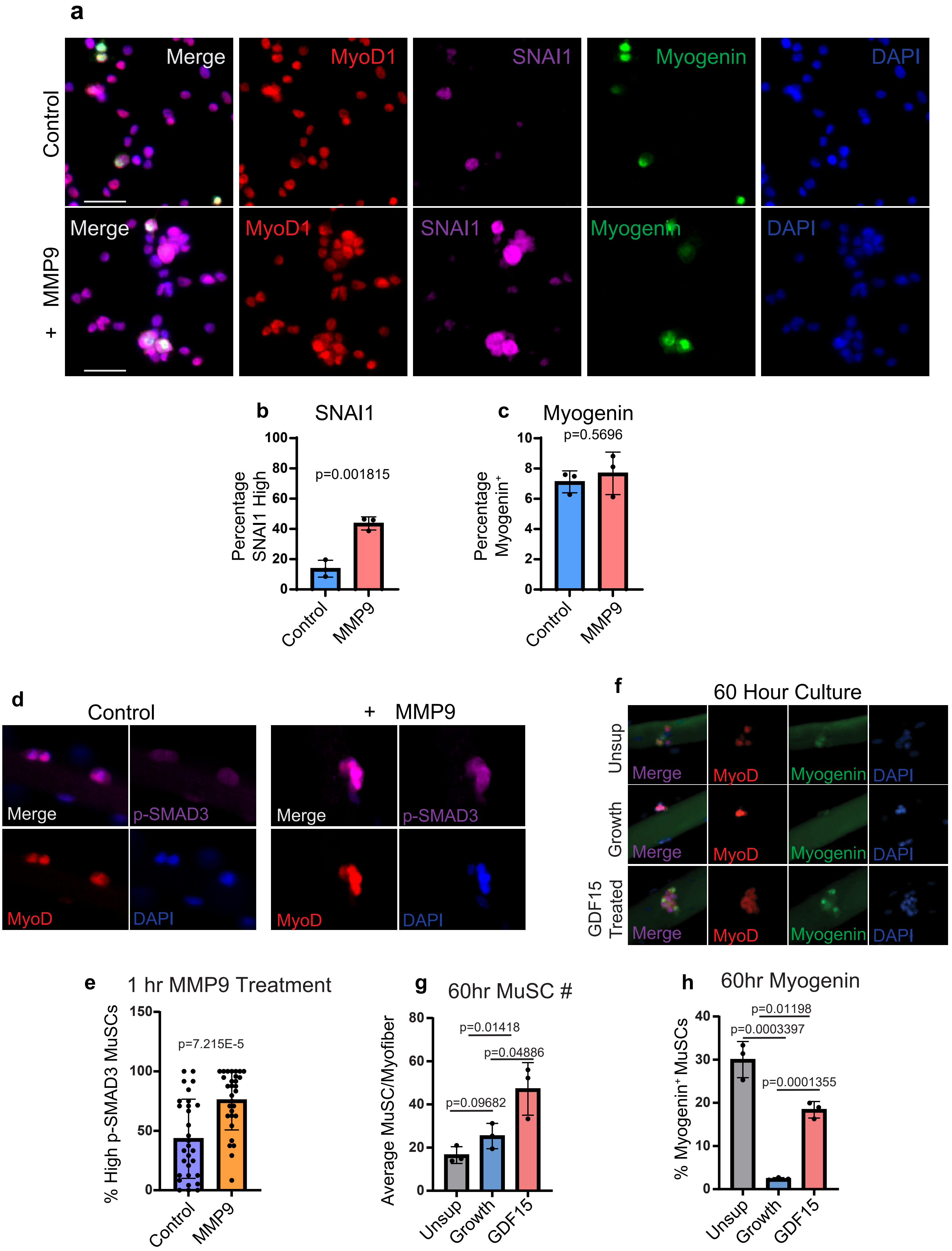

**Extended Data 9: Primary myoblasts cocultured with GDF15 KO C26 cells show no phenotypic improvement**

**a.** Representative images of a 12-hour EDU incorporation assay performed on Crispr Control and GDF15 KO C26 cells. **b.** Quantification of the percentage of EDU positive C26 hours after 12-hour EDU incorporation. **c.** Representative images of a 12-hour EDU incorporation assay performed on primary myoblasts that were not cocultured, or cocultured with Crispr Control or GDF15 KO C26 cells. **d.** Quantification of the percentage of EDU positive primary myoblasts after a 12-hour EDU incorporation. **e.** Representative images of primary myotubes that were differentiated for 5 days and were either not cocultured, or cocultured with Crispr Control or GDF15 KO C26 cells. **f.** Bar graph of the fusion index of primary myotubes that were differentiated for 5 days and were either not cocultured, or cocultured with Crispr Control or GDF15 KO C26 cells. **g.** Representative images of isolated EDL myofibers that were either not cocultured, or cocultured with Crispr Control or GDF15 KO C26 cells for 72 hours. Myofibers were stained for MyoD and Myogenin. Scale bar = 40 µm. n = 3 biological replicates. **h.** Quantification of the number of MuSCs per myofiber on isolated EDL myofibers cultured for 72 hours in growth media, or cocultured with Crispr Control or GDF15 KO C26 cells. **i.** Percentage of Myogenin positive MuSCs per myofiber in isolated EDL myofibers cultured for 72 hours in growth media, or cocultured with Crispr Control or GDF15 KO C26 cells.

Extended Data 9

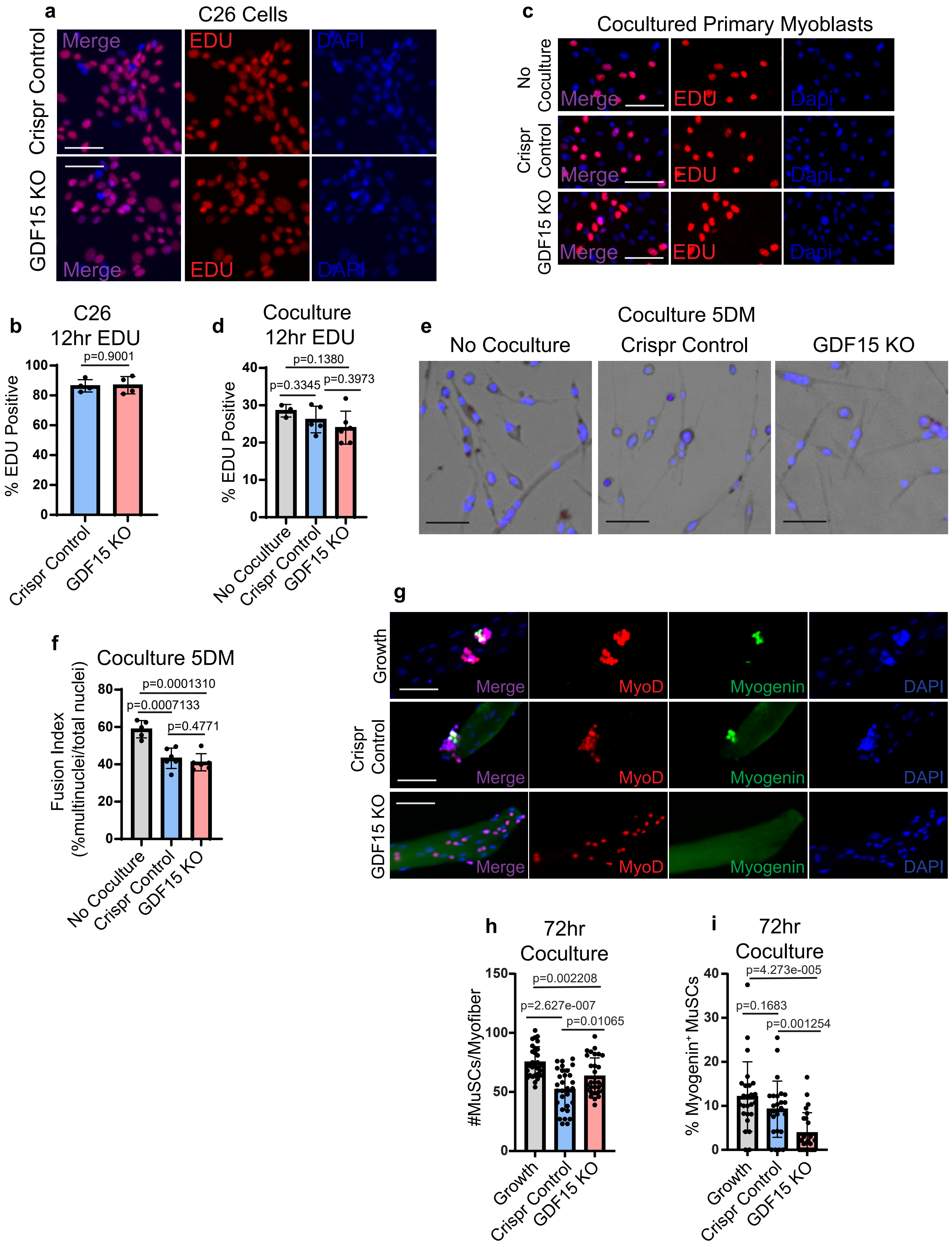
